# Endosymbiosis: metabolic trade-offs drive reproductive synchronization

**DOI:** 10.1101/2025.10.19.683235

**Authors:** Lucas Santana Souza, Eric Libby

**Affiliations:** Department of Mathematics and Mathematical Statistics, Umeå University, Umeå, Sweden; Integrated Science Lab, Umeå University, Umeå, Sweden; Umeå Center for Microbial Research (UCMR), Umeå University, Umeå, Sweden

**Keywords:** endosymbiosis, synchronization, metabolism, mathematical modeling

## Abstract

Endosymbiotic relationships have fueled the evolution of complex life. The persistence of these relationships relies on coordinating the reproduction of hosts and their guests. Yet, such coordination may come at a cost, as one or both partners may need to slow their growth to synchronize. Here, we examine the fitness consequences of synchronization by using a combination of mathematical and bacterial metabolic models. Analyzing millions of putative host–guest pairings, we find that synchronization is typically costly for hosts but beneficial for guests. We show that synchronization can occur when hosts relinquish metabolic resources to guests. Given the costs to hosts, we investigate whether hosts can benefit from growing faster, at the risk of losing their endosymbionts. Our mathematical model suggests this outcome is theoretically possible, but metabolic modeling consistently indicates that synchronization increases overall fitness— provided the formation of new endosymbioses is rare. When new relationships can readily form, the optimal strategy shifts: hosts maximize fitness by growing as fast as possible, leading to regular cycles of loss and reacquisition of their guests. Overall, our findings indicate that while synchronization is costly for hosts, it offers long-term fitness benefits in contexts where endosymbionts are not easily replaced.

## Introduction

Endosymbioses play a pivotal role in the evolution of complex life (1–4). They provide species with new functional capacities and drive the formation of new internal structures, such as organs and organelles (5–8). Yet, in order to realize this evolutionary potential, endosymbioses must persist, and this is often achieved via synchronized reproduction between species. Without synchronization, the mismatch in reproductive rates could destabilize the relationship (9–12). If guests proliferate faster than their hosts, they may overuse the host’s resources, like a cancer, and harm the host (13). Conversely, if hosts replicate faster than their guests, the host population could eventually lose all of its guests (14–17). While synchronizing growth rates can help maintain an endosymbiosis, it often requires one of the species to sacrifice a possibly higher individual growth rate in service of the pair (17, 18). If this sacrifice is too costly, then it could cause the endosymbiosis to dissociate, returning host and guest to their free-living lifestyles (18, 19). Here, we investigate whether synchronization offers sufficient benefits to outweigh its potential costs.

One benefit of synchronization is that it can act to preserve the symbiotic partnership across future generations, i.e. it ensures successful vertical transmission. If hosts reproduce faster than guests, then eventually a host will reproduce and its offspring will be without an endosymbiont (guest). Assuming there is some benefit to the endosymbiosis— hosts with guests are fitter than those without— there is selective pressure for hosts to maintain their guests, especially after reproduction. Even in established endosymbioses where synchronization is coordinated by complex regulation, there are still opportunities for missteps that cause hosts to lose guests. For example, yeast can lose their mitochondrial DNA, forming so-called petite colonies, even though the mitochondria are an organelle derived from an endosymbiont present in early eukaryotes (20–22). The threat of failed vertical transmission is likely to be more pronounced in early endosymbioses that have yet to evolve robust regulatory mechanisms. Indeed, vertical transmission is guaranteed to fail if the guest does not reproduce before the host. Thus, in the context of early endosymbioses, synchronization of growth rates is a likely first step in the process of establishing a successful long-term partnership.

Another benefit of synchronization is that it can act to reduce the variance of the number of guests in hosts. Suppose hosts with a specific number of guests have the highest fitness. Maximizing the host’s long-term fitness equates to ensuring that hosts reliably maintain the same number—the optimal number— of guests. This same argument appears in other evolutionary contexts that consider populations with variable fitness, e.g. populations with bet-hedging or phenotypic plasticity (23–27). In these cases, geometric mean fitness typically determines the long-term success of genotypes, and optimizing geometric mean fitness is done by reducing variance (28–33). By synchronizing reproduction, hosts can more reliably maintain the same number of guests across generations.

The cost of synchronization depends on which partner would grow faster if not synchronized. If hosts outpace their guests, they must either slow their own reproduction or create conditions that help guests catch up. Either way, hosts sacrifice potential growth in order to synchronize. If, instead, guests outpace their host, the challenge reverses. Rather than hosts slowing their growth, they must find ways to limit guest proliferation, possibly by restricting access to nutrients or other resources (11, 34–36). Yet imposing such limits may come with its own costs in terms of regulation and enforcement (35, 37, 38). Thus, whether hosts are slowing down or holding guests back, synchronization requires a reallocation of nutrients and energy, which can ultimately translate to a fitness cost (39).

Given the various benefits and costs of synchronization, it can be difficult to determine which should be expected to dominate. In order to properly evaluate the net effect of synchronization, we would need to know the growth rates of hosts and guests across different scenarios: when they synchronize, when they do not synchronize (here, called “uncoordinated”), and when they live independently, i.e. not in an endosymbiosis. Such detailed information may be possible to acquire in specific empirical contexts, but would be challenging to get more widely. To circumvent these challenges, we use genome-scale metabolic models. These models utilize genomic information to infer the metabolic repertoires of organisms and predict their growth rates in various ecological and environmental contexts (40, 41). They have also been used to address more general evolutionary questions (42, 43), including—and of relevance here— endosymbioses (44). One advantage of genome-scale metabolic models is the availability of large collections of models, particularly for prokaryotes. By leveraging these large collections, we can analyze a wide range of endosymbioses and extract general patterns concerning when the benefits of synchronization outweigh its costs.

In this study, we quantify the benefits and costs of synchronization between hosts and guests engaged in metabolic endosymbioses. To do this, we first use genome-scale metabolic models to construct a large set of hypothetical endosymbioses between pairs of bacteria (31 million pairs). We then estimate the growth rates of hosts and guests in each endosymbiosis in three different possible scenarios: synchronized growth, uncoordinated growth, and free-living growth. By comparing growth rates between these scenarios, we identify how often the host or guest pays a cost for synchronization. We then use the computed growth rates as parameters in a dynamical model and solve for when synchronization improves a host’s fitness (growth rate) compared to uncoordinated growth. Lastly, we consider the extent to which establishing new endosymbioses (i.e. horizontal transmission) alters the relative costs and benefits of synchronization. Our results reveal that synchronization is favored in the vast majority of cases, provided that forming endosymbioses is exceptionally rare; otherwise, it can be advantageous for hosts to not synchronize with their guests and thereby rely on regularly, re-occurring transient endosymbioses.

## Results

We start by investigating whether synchronized growth in endosymbioses affects their initial establishment. In synchronized growth, resources must be allocated so that hosts and guests have the same growth rate. Previous work exploring the rarity of bacteria-bacteria endosymbioses (44) assessed their possible viability by assuming synchronization; however, this assumption could exclude cases of viable endosymbioses where hosts and guests cannot synchronize, i.e. they can only grow at different rates (uncoordinated growth). To determine the growth rates of hosts and guests under synchronized and uncoordinated growth scenarios, we use genome-scale metabolic models from two of the largest collections: AGORA (818 models) and CarveMe (5,587 models) (see *“Structure of genome-scale metabolic models”* in Methods for details). Fig. 1A depicts how we use these models to determine viability in endosymbioses with synchronized/uncoordinated growth (see *“Growth rate calculations”* in Methods for details). We note that in uncoordinated growth, we assume the host has primary access to resources to maximize its growth, and then the guest maximizes its growth on the remaining resources. A consequence of this assumption is that there is a possibility that the guest cannot grow because an essential resource was consumed by the host. Thus, it is unclear whether synchronized or uncoordinated growth is more likely to restrict viable endosymbioses.

**Fig. 1.**
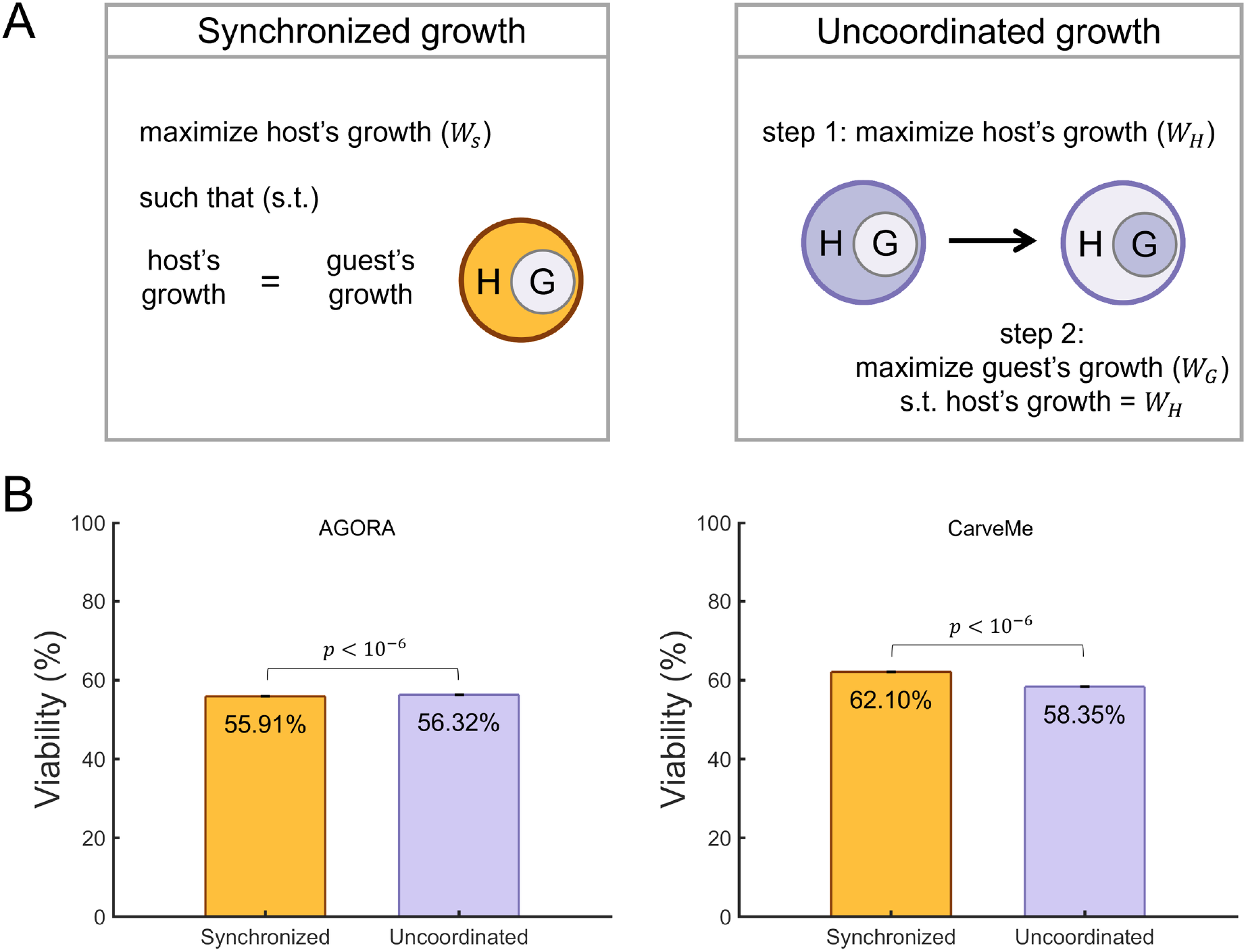
Viability of endosymbioses with synchronized and uncoordinated growth. (A) A schematic illustrates the computation of the synchronized and uncoordinated growth rate of a putative endosymbiosis using metabolic models. For synchronized growth, we use flux balance analysis to maximize the host’s growth rate, while requiring that the growth rates of the host and its guest are equal (see *“Growth rate calculations”* in Methods). For uncoordinated growth, we first maximize the host’s growth rate and then maximize the guest’s growth, requiring that the host still grows at its maximum rate. (B) The bar graph shows the percentages of paired host-guest metabolisms that form viable endosymbioses from AGORA and CarveMe under either synchronized growth (orange) or uncoordinated growth (purple). The percentages are the mean, and the error bars show the standard error of a total of 669,124 pairs for AGORA and 31,214,569 pairs for CarveMe. For AGORA, the difference between synchronized and uncoordinated growth scenarios is small (≈ 0.41%) though statistically significant (McNemar’s test, *p* < 10^−6^). For CarveMe, there is also a small difference in the viability between synchronized and uncoordinated growth scenarios (≈ 3.75%), which is bigger than in AGORA and also statistically significant (McNemar’s test, *p* < 10^−6^).

In Fig. 1B, we show the percentage of viable putative endosymbioses for all possible pairings of metabolic models within AGORA (669,124 pairs) and CarveMe (31,214,569 pairs). Overall, we find similar levels of viability between synchronized and uncoordinated growth scenarios, with differences of 0.41% for AGORA and 3.75% for CarveMe. While small, the differences are statistically significant and show opposite trends. In AGORA, the uncoordinated scenario has the highest viability (McNemar’s test, *p* < 2.2 × 10^−16^). In CarveMe, the synchronized scenario has the highest viability (McNemar’s test, *p* < 2.2 × 10^−16^). Furthermore, we observe that approximately 0.7% of pairs in AGORA (and 0.2% in CarveMe) that are unviable under synchronized growth become viable under uncoordinated growth (see Tables S1-S2). Conversely, around 0.3% of pairs in AGORA (and 4% in CarveMe) that are unviable under uncoordinated growth become viable under synchronized growth (see Tables S1-S2).

After establishing that synchronization does not have a large effect on viability, we turn our attention to its impact on growth rate. We consider all endosymbioses that were viable in both uncoordinated and synchronized growth scenarios from Fig. 1 (372,120 pairs in AGORA and 18,147,674 pairs in CarveMe) and identify how synchronization affects the growth of the host and guest. Fig. 2A shows the possible types of effects we may observe, organized into three categories: costly (the growth rate is lower in the synchronized scenario), neutral (the same), or beneficial (higher). We note that because of our methodology, a host’s growth rate can never be faster in the synchronized case; otherwise, it would have grown at this rate in the uncoordinated case as well. Thus, we are left with six possible outcomes (see Fig. 2A). Fig. 2B shows that the most common outcome that occurs in *>* 99% of pairs is that synchronization is costly for the host but beneficial to the guest. This occurs because in the uncoordinated scenario, hosts grow as fast as possible, leaving few resources for guest growth, and so hosts grow faster than their guests. The second most common outcome–though far rarer (0.6% in AGORA or 0.38% in CarveMe)–is that synchronization is beneficial for the guest but has no effect on the host, which means that synchronization imposes no burden on the host. We also observed instances where synchronization was costly for both hosts and guests. In these cases, synchronization is achieved by forgoing a set of resources and their products that benefit both partners, though especially the host. Finally, we identified instances where the growth rates under the uncoordinated scenario naturally led to synchronization, though this was rare: fewer than 0.01% of pairs (6 pairs) in AGORA and fewer than 0.001% (178 pairs) in CarveMe (see species names at Tables S4-S5).

**Fig. 2.**
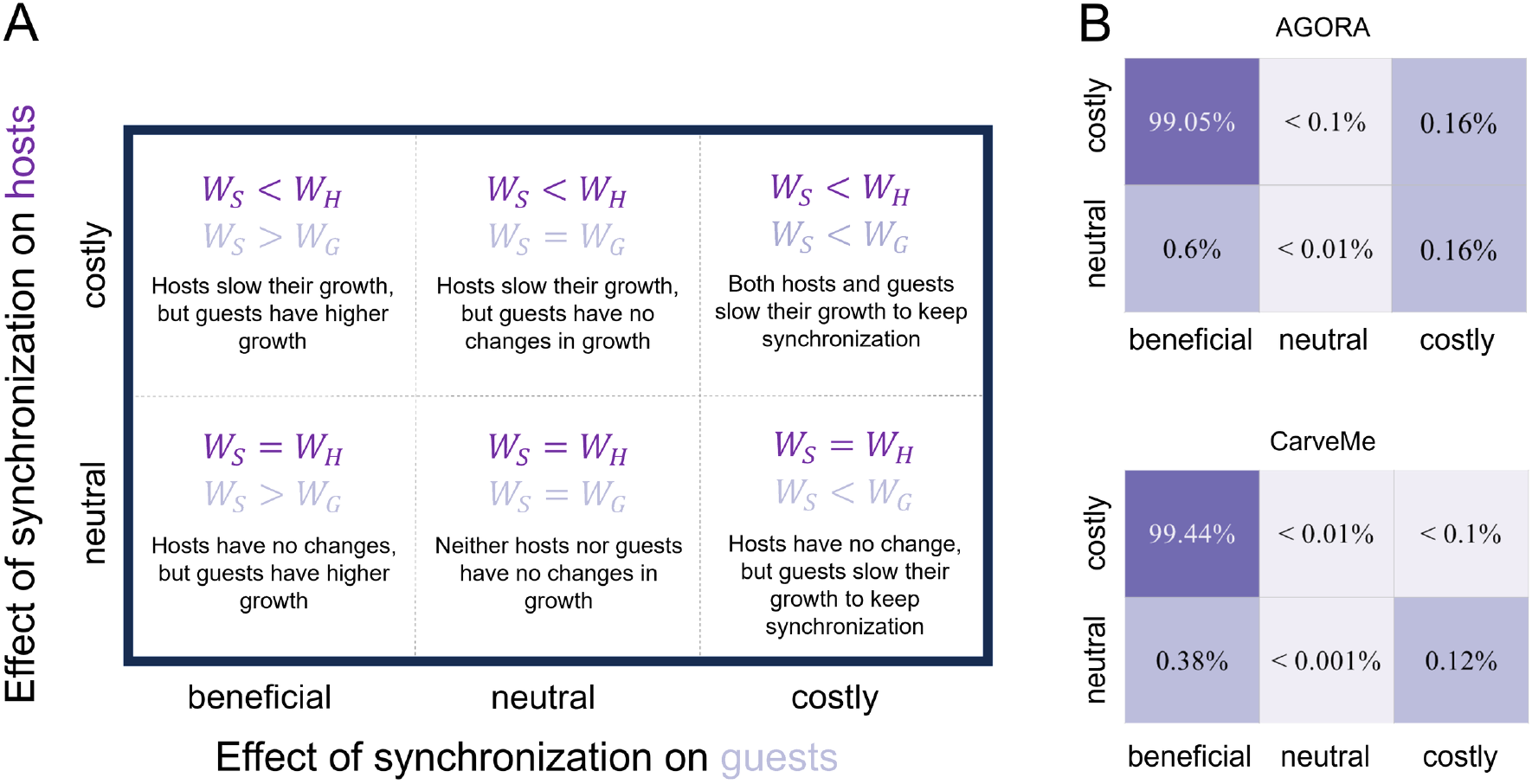
The impact of synchronization on the growth rate of hosts and guests. (A) The schematic categorizes the possible effects of synchronization on the growth rate of hosts (rows) and guests (columns) using our metabolic methodology. The effects are either costly (the growth rate is lower in the synchronized scenario), neutral (the same), or beneficial (higher). We note that hosts cannot increase their growth rate by synchronizing with their guests, i.e. synchronization is never beneficial for hosts. (B) The heatmap shows the percentages of host-guest pairs that fall into each of the six outcomes from (A) for AGORA and CarveMe. These percentages were calculated from all viable pair combinations under the synchronized and uncoordinated scenarios (372,120 pairs for AGORA and 18,147,674 pairs for CarveMe). For both collections, the most common outcome (> 99% of pairs) is that synchronization is costly for the host but beneficial to the guest, which means in the uncoordinated scenario that hosts grow faster than their guests.

Given that in the majority of cases synchronization benefits guests while imposing a cost on hosts, this suggests a possible pathway from uncoordinated growth to synchronization. If hosts forgo using some resources so that their guests can use them (or if hosts use other pathways that create resources for guests), it could simultaneously lower the growth rate of hosts and raise the growth rate of guests. We explored this possible path to synchronization in 100 randomly selected host-guest pairs (see species names at Tables S6-S7) by starting from an initial uncoordinated scenario and then iteratively requiring hosts to increase the growth of their guests (see Fig. 3A and *“Path to synchronous growth”* in Methods). In all considered cases, we were able to trace a path from uncoordinated growth to synchronization with similar qualitative features in AGORA and CarveMe models (see Figs. 3B-D). Fig. 3B shows that in the uncoordinated state, hosts grow an average of 40% − 60% faster than in the synchronized state, while the guest’s growth rate is near 0. As hosts allocate resources to guests, the growth rates of guests increase sharply. For example, Fig. 3D shows that sacrificing just 1% of host growth can lead to a more than 100-fold increase in guest growth in over 80% of tested pairs. This trend persists across different levels of synchronization, measured as the guestto-host growth rate ratio (*W*_*G*_/*W*_*H*_). As shown in Fig. 3C, the average net gain in guest growth consistently exceeds the net loss in host growth, indicating that sacrifices by the host yield larger benefits for the guest (for the individual plot of each pair, see Fig. S1).

**Fig. 3.**
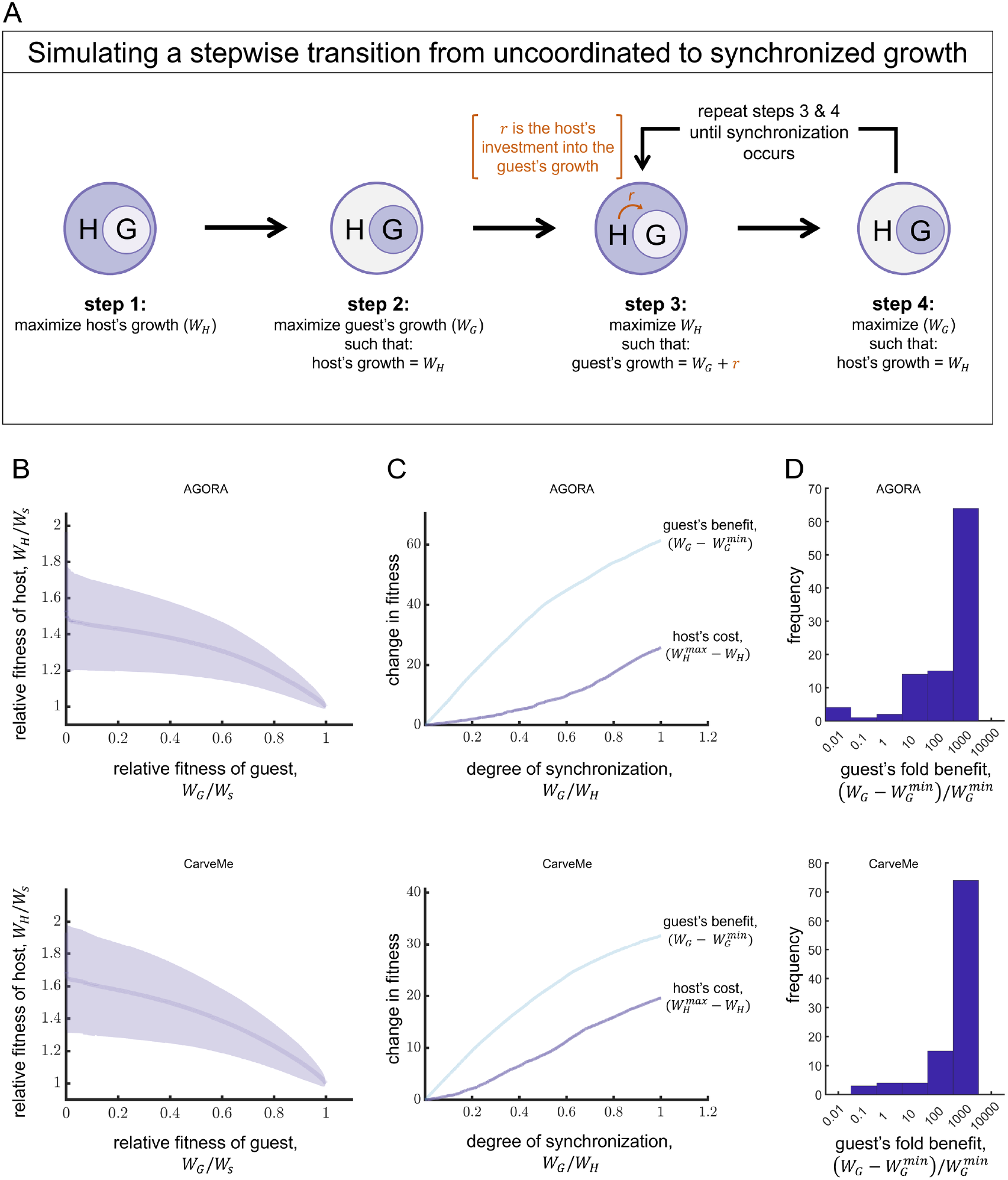
Fitness trade-offs in reproductive synchronization. (A) A schematic illustrates the method used to calculate growth rates (fitness) across a transition from uncoordinated to synchronized growth (see *“Path to synchronous growth”* in Methods). Steps 1-2 calculate the growth rate of hosts and guests under the uncoordinated scenario. At these steps, the host’s fitness is the highest 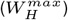, and the guest’s fitness is the lowest 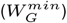. Steps 3-4 calculate the fitness of hosts (*W*_*H*_) and guests (*W*_*G*_) until synchronization is reached and both have the same fitness (*W*_*S*_). (B) Host’s fitness relative to synchronization (*W*_*H*_ /*W*_*S*_) is plotted against guest’s fitness relative to synchronization (*W*_*G*_/*W*_*S*_). The dark line and shaded area represent the mean and standard deviation in *W*_*H*_ /*W*_*S*_ across 100 distinct pairs from AGORA (top) and CarveMe (bottom). The leftmost values show that hosts grow an average of 40% (AGORA) or 60% (CarveMe) faster in the uncoordinated scenario compared to synchronization. From left to right, the graphs show how fitness changes as the pair becomes more synchronized; in particular, early increases in synchronization incur low costs for hosts (as indicated by the flat slope). (C) Plotted is the average fitness benefit for guests and fitness cost for hosts as a function of the degree of synchronization, measured as the ratio of their growth rates (*W*_*G*_/*W*_*H*_). The benefit for guests is the average change in fitness relative to its minimum value when there is no synchronization 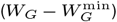. The cost for hosts is the average change in fitness relative to its maximum value in the absence of synchronization 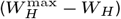. For all degrees of synchronization, the average guest fitness benefit exceeds the fitness cost experienced by hosts. (D) The histogram shows the distribution of guest’s fold benefit at the point where the host has lost 1% of its maximal fitness, i.e. 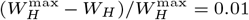. Incurring this minor cost to the host results in more than a 100-fold increase in guest growth for over 80% of tested pairs.

The results of Fig. 3 show that there is a range of intermediate growth rates (for hosts and guests) between the uncoordinated scenario and synchronization. In each of these states, the host experiences a tradeoff between its growth and the longevity of the partnership. To determine when synchronization of an endosymbiosis may be favored, we developed a mathematical model linking host population dynamics with resource availability (see Fig. 4A for a schematic). We consider three host populations: hosts that reproduce in a synchronized manner with their guest (*H*_*s*_); hosts that reproduce without coordinating with their guest (*H*_*u*_); and hosts that have lost their guest (*H*_0_). Each type of host (*H*_*s*_, *H*_*u*_, and *H*_0_) consumes the same common resource *R* at a rate *c* and grows at a maximum rate *µ*_*s*_, *µ*_*u*_, and *µ*_0_, respectively, following Monod kinetics with a half saturation parameter *θ*. The resource flows into the environment at a rate *k*_*r*_, and the resource and all hosts experience an outflux rate *ϕ* (Table S3). Since hosts and guests reproduce at different rates in the uncoordinated case, there is a rate *d* at which the partnership dissociates. Thus, when a *H*_*u*_ host reproduces, it gives rise to an *H*_0_ host. For simplicity, we assume that in the synchronized case, hosts and guests never dissociate. Finally, we assume that the formation of new endosymbioses is so rare as to be negligible, though later we consider when this assumption is violated. The resulting dynamics of the resource and host populations are described by the following set of differential equations:

**Fig. 4.**
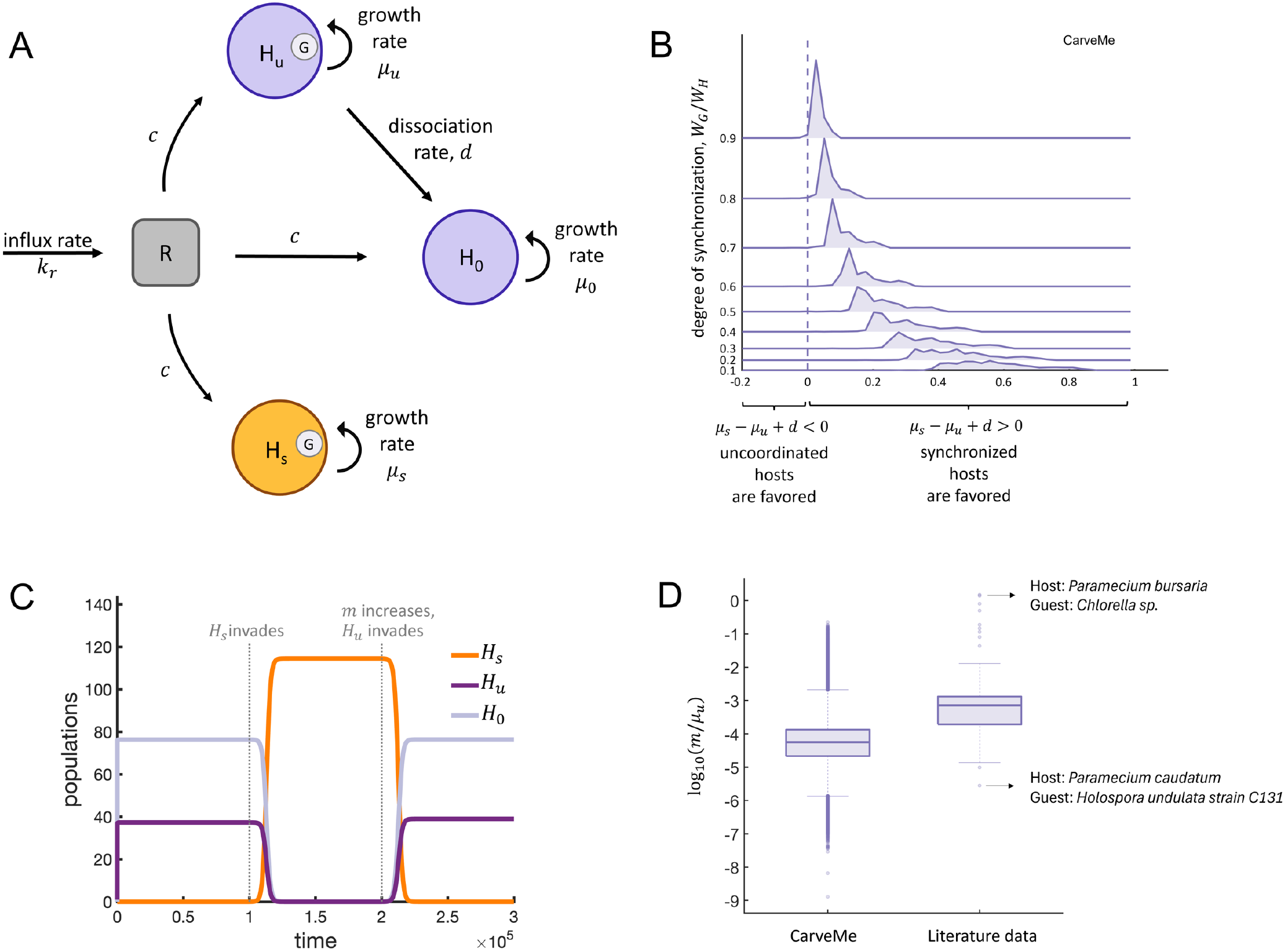
Mathematical model of the competition between synchronization and uncoordinated growth. (A) A schematic of the mathematical model in Eq. (1) shows the various host populations (*H*_*u*_, *H*_*s*_, *H*_0_) competing for a limiting resource *R*. (B) A joyplot displays a series of histograms for different degrees of synchronization (*W*_*G*_/*W*_*H*_). Each histogram shows the distribution of the differences in net growth rate between synchronized (*µ*_*s*_) and uncoordinated growth (*µ*_*u*_ − *d*) in 1000 host-guest pairs using CarveMe models (the corresponding graph for AGORA is Fig. S3A). The dashed line separates where uncoordinated growth (left) and synchronization (right) are favored. In all host-guest pairs studied, synchronization results in higher growth. (C) A numerical solution of the mathematical model from A) shows the population dynamics of host populations in response to invasion and the formation of new endosymbioses. Using the parameters (*c* = 0.1, *k*_*r*_ = 1, *θ* = 5, *ϕ* = 0.1, *m* = 0.0038), the population starts with only an uncoordinated host (*H*_*u*_) and quickly reaches a steady state with a mixed population of *H*_*u*_ and the host alone (*H*_0_). Although *H*_*u*_ reproduces faster than *H*_0_ — even when accounting for dissociation (*µ*_*u*_ = 2.4, *µ*_0_ = 0.6, *d* = 1.3)— the *H*_0_ population reaches a higher proportion of the population. At *t* = 10^5^, a synchronized host (*H*_*s*_) invades at a small fraction and displaces the *H*_*u*_ and *H*_0_ populations, because its growth rate (*µ*_*s*_ = 1.2) is higher than *µ*_*u*_ −*d*, and the rate that new endosymbioses form is below the threshold (*m*\* = 0.0042). At *t* = 2 × 10^5^, the rate that new endosymbioses form is raised so that it is above the threshold (*m* = 0.0046 > 0.0042). This allows the uncoordinated host to be able to invade from rare and drive the synchronized host to extinction. (D) Boxplots show the distribution of the endosymbioses formation rate (*m*) scaled by uncoordinated host growth rate from two different datasets: 1. our modeling analyses (labeled CarveMe) and 2. data in the literature. For the CarveMe boxplot, we used the critical rate at which synchronization is disfavored *m*\* (Eq. (5)) as the value of *m*, with *ϕ* = 0.01. For the literature data boxplot, we calculated *m* as described in *SI-B* and used *µ*_*u*_ = *ln*2/24, where 24 denotes the doubling time (in hours) of the population. In each box, the central mark indicates the median, the bottom and top edges represent the 25th and 75th percentiles, and the whiskers extend to the most extreme data points not considered outliers, which are plotted as individual dots. Values from the experimental observations are higher on average than those from our modeling analyses. The CarveMe boxplot uses 2,452,637 measurements (all distinct host–guest pairs where synchronizing hosts are fitter when *m* = 0) and the literature data boxplot uses 78 measurements. The corresponding graph for AGORA is found in Fig. S3B.

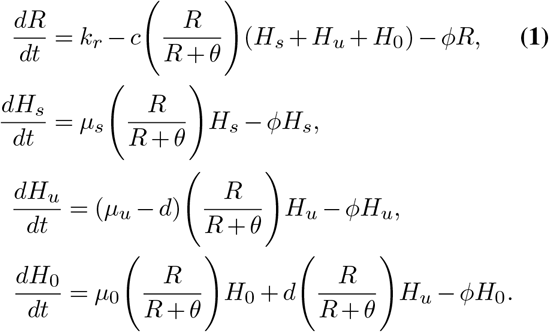

Since we are interested in when synchronization with a guest may confer an advantage, we assume that hosts with a guest grow faster than hosts without one, i.e *µ*_*s*_ > *µ*_0_ and *µ*_*u*_ > *µ*_0_. If this were not true, then it would always be better to expel the guest than synchronize reproductive rates. We also assume that uncoordinated endosymbioses grow faster than synchronized ones, i.e. *µ*_*u*_ > *µ*_*s*_. If this were not true, then it would always be best to synchronize reproductive rates. Moreover, our metabolic modeling results indicate that this is the most common scenario. Based on the ordering of the growth rates—and the implicit assumption that the influx of resources is high enough to sustain populations—we find that there are two possible steady states: 1. the population consists of only synchronized host-guest pairs, and 2. there is a mix of uncoordinated host-guest pairs and hosts without guests. In the second steady state, hosts without guests (*H*_0_) persist because they are continually produced by the dissociation of uncoordinated endosymbioses. A stability analysis of the mathematical model indicates that which steady state is reached depends on the relative values of *µ*_*u*_ − *d* and *µ*_*s*_, such that synchronization is favored when *µ*_*s*_ > *µ*_*u*_ − *d* or equivalently when 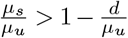.

We can determine whether synchronization is more often favored than uncoordinated growth by using metabolic modeling to estimate the relative values of *µ*_*u*_, *µ*_*s*_, and *d*. The growth rates *µ*_*u*_ and *µ*_*s*_ follow immediately as they are proportional to the growth rates (*W*_*H*_ and *W*_*S*_) calculated by flux balance analysis, and so 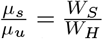. The dissociation rate *d* is not as direct, but it can be derived from the growth rates of the host and guest in an endosymbiosis. The full derivation is in the Supplementary section (*“Derivation of dissociation rate”*), but we present a brief description here. To estimate *d*, we consider host reproduction in terms of discrete reproductive events. For the host and guest to stay together, the guest must replicate by the time the host does; otherwise, when the host divides, one offspring will be without a guest. So, in some interval of time, the number of reproductive events for hosts and guests can be used to estimate the rate of dissociation. In particular, it is the ratio of reproduction times that determines the frequency of dissociation events. By relating this ratio of reproduction times to the ratio of continuous growth rates, we can derive the dissociation rate *d*:

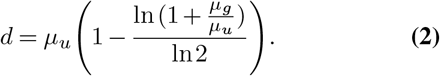

The *µ*_*g*_ term is the growth rate of the guest in the endosymbiosis, but it only appears as a ratio with the host’s growth rate. Using this expression for *d* and the equivalence of the growth ratios between mathematical and metabolic models, we find that synchronization is favored over uncoordinated growth when:

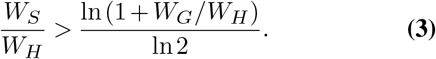

We can ascertain when Eq. (3) holds by using the calculations from the tradeoff curves in Fig. 3B. Fig. 4B shows that for all of the pairs we have considered, synchronization is fitter than uncoordinated growth. Synchronization is also fitter compared to the intermediate cases when hosts act to increase the growth of their guests. A quick calculation can give some insight into why this is the case. At the extreme, when the host does not limit its own growth, the relative advantage for the host compared to synchronized growth (*W*_*H*_/*W*_*S*_) is typically less than 3 (see Fig. S2). Substituting this value into Eq. (3) shows that synchronization is favored when the value of *W*_*G*_/*W*_*H*_ is less than 25%. The ratio *W*_*G*_/*W*_*H*_ corresponds to the growth of the guest relative to the host, and at the extreme of uncoordinated growth, its value is closer to 1%, i.e. it is significantly below 25% (see Fig. 3C). Values of the ratio *W*_*G*_/*W*_*H*_ that are closer to 25% correspond to when hosts relinquish resources to guests and are consequently growing slower relative to the uncoordinated extreme. This slower growth has its own critical value of *W*_*G*_/*W*_*H*_ that marks when it is better to synchronize, and it is higher than 25%, thereby creating a moving target.

Although we have found that synchronization increases fitness, we have made the restrictive assumption that once the guest is lost, it cannot be regained. We relax this assumption by assuming that there is a constant rate *m* that new endosymbioses form. The equations for the dynamics of *H*_*u*_ and *H*_0_ then become

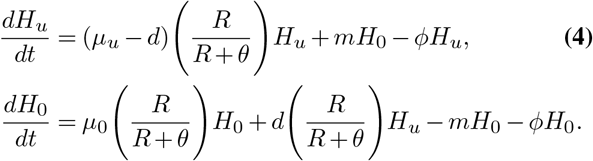

With this change, we expect that high values of *m* should favor uncoordinated growth and low values of *m* should favor synchronization. We calculate the critical value of *m* (denoted as *m*\*) for which synchronization is no longer favored and find:

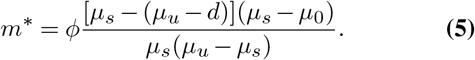

In Fig. 4C, a numerical solution of the differential equations model shows that when *m* increases past *m*\* a population of synchronized hosts can be invaded and replaced by hosts with uncoordinated growth.

While the value of *m*\* in Eq. (5) depends on five different terms, we focus on one in particular, *d*. Since the dissociation rate *d* describes how often endosymbioses dissolve, we would expect the critical value of their formation, i.e. *m*\*, to be proportional, so that higher values of *d* require higher values of *m*\* to favor uncoordinated growth. We calculated *m*\* for all host–guest pairs (2,452,637 in CarveMe and 53,591 in AGORA) where synchronization is the optimal strategy when new endosymbioses do not form, i.e. when *m* = 0. We rescaled *d* and *m*\* by growth rates so that they could be compared using data from the metabolic models (*see SI– “Comparison of m*\* *and d”*). We then used these values to calculate the coefficient of determination (details at *SI–*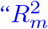 *calculation”*). Counter to our initial expectation, we found only a very weak association: *d* explained less than 2% of the variance in *m*\* in both AGORA and CarveMe collections (Fig. S4).

Having calculated *m*\*, we also asked whether these values fall within biologically realistic ranges. To evaluate this, we estimated *m* from published empirical observations (45– 51) (details in *SI – “m estimation”*). This dataset encompasses 43 distinct host–guest pairs and 78 measurements (Table S8). We note that in this dataset endosymbioses readily form between free-living hosts and guests. Using these data, we found that all of our scaled *m*\* estimates fell below the range observed empirically (Fig. 4D and Fig. S3B). Thus, the critical formation rate *m*\* for endosymbioses appears to lie within biologically plausible bounds.

## Discussion

A hallmark of many established endosymbioses is reproductive synchronization. While synchronization can act to maintain an endosymbiotic relationship across generations, it also imposes potential costs by requiring at least one partner species to slow its growth. In this study, we combined mathematical modeling and metabolic analyses to weigh the costs and benefits of growth synchronization in simulated prokaryotic endosymbioses. We found that when the formation of new endosymbioses is exceptionally rare, synchronization is favored in the vast majority of cases. We also discovered that if hosts forego certain metabolic resources, they can increase the growth rate of their guests and thereby increase their own fitness by prolonging the partnership. Thus, we identified a plausible evolutionary route by which initially uncoordinated growth could transition to synchronization. A crucial factor shaping these outcomes was the possibility of reforming endosymbioses. Even at low frequencies, the ability to re-establish partnerships reduced the advantage of synchronization and instead favored a dynamic life cycle in which uncoordinated hosts reproduce quickly, lose their guests, and reacquire them from the environment.

A main result from our analyses is that the synchronization of growth rates in an endosymbiosis is overwhelmingly favored in comparison with uncoordinated growth. Using metabolic models to compute growth rates, we found that hosts could grow up to 3 to 4 times faster—though 99% are less than 2 (see Fig. S2)— by exploiting their guests rather than synchronizing. Yet, this growth benefit is short-lived because it creates a disparity between host and guest growth rates that prevents the endosymbiosis from reliably reproducing across generations. If we assume that the endosymbiosis confers some fitness benefit, then loss of a guest poses an evolutionary cost. We analyzed a mathematical model comparing uncoordinated and synchronized endosymbioses and solved for the critical condition that determines which is fittest. Our analytical solution shows that the dissociation rate plays a key role in the competitive outcome of endosymbioses. Further metabolic modeling revealed that hosts could forego resources to increase the growth rate of their guests, thereby lowering the dissociation rate. We characterized the tradeoffs between growth and the stability of the endosymbiosis along the spectrum from uncoordinated growth to synchronization (Fig. 4B and Fig. S3A). In all the studied cases, synchronization resulted in higher fitness for the entire spectrum except when the difference between host and guest growth rates was small, i.e. close to the synchronized rate. Thus, even though synchronization comes at a cost in terms of host growth, the increased stability of the endosymbiosis more than compensates.

The benefits of synchronization were sensitive to one assumption: that free-living hosts and guests formed new endosymbioses so rarely that they could be ignored. When we relaxed this assumption, we found that uncoordinated growth could be favored over synchronization provided that new endosymbioses formed above some critical rate *m*\* (Fig. 4C). This critical rate *m*\* was specific to host-guest pairs and only had a weak relationship with the rate at which the endosymbiosis breaks apart. We calculated this critical rate for different host-guest pairs using metabolic models and compared it with empirical data from systems where new endosymbioses can be observed forming. We found that the rates required by our analytical models were well within the range of biological observations, suggesting that the environmental acquisition of guests could present a major impediment to the evolution of synchronization.

The results from our modeling reflect the fact that extant endosymbioses exhibit both synchronized and uncoordinated growth (15–17). A canonical example of synchronization occurs in cells where guests have become organelles, or are on an evolutionary trajectory toward organelle status. In contrast, cases of uncoordinated growth can sometimes be difficult to distinguish from failures of synchronization, as seen in yeast petite colonies that lose their functional mitochondria. One extreme example of true uncoordinated growth occurs in *Hatena arenicola* where, upon host division, only one daughter cell inherits the Nephroselmis guest, while the other daughter cell must acquire a new guest from the environment (52, 53). This system resembles the cases of uncoordinated growth we explore, where the host gains a benefit from the guest, even though the guest effectively does not reproduce within the host. Given the diversity of host-guest interactions, we expect that different selective pressures are likely shaping the degree of synchronization. Mathematical models have identified mechanisms that favor synchronization (10, 16, 54), such as obligate host-guest dependencies (55) or enhanced survival in resource-poor environments (56). Our work shows how metabolism and selection for fast growth can drive synchronization through resource sharing, or alternatively, maintain cyclic gains and losses of endosymbioses due to uncoordinated growth.

In our metabolic analyses, we assumed that hosts managed the resource pools inside the cell. Based on this assumption, hosts could exert control over guests by consuming resources themselves or by leaving them available for guest use. In this framework, synchronization depends primarily on host actions, and whether it is favored depends on how these actions affect host fitness across different scenarios. There are many examples of endosymbioses where hosts are thought to exert control (11, 12, 17, 35, 36, 57–60), using diverse mechanisms such as gene regulation or the expulsion/destruction of guests (12). However, there are also cases where the guest exerts some level of control over host reproduction (17, 50, 61–64). For example, in (65), the authors consider a case in which the guest limits its own growth. In our system, we could simulate the guest having control by allowing it priority access to resources. While we expect that the structure of our mathematical models would remain similar (apart from a focus on guest growth rates), the growth parameters would differ. A previous study found that in the vast majority of prokaryotic endosymbioses, free-living guests grow faster than those in synchronized endosymbioses, because within a host, their growth is constrained by the resources the host can import (44).

We note that our results are based on a particular type of endosymbiosis: a metabolic interaction between two prokaryotic species. We chose this scenario for several reasons. First, we focused on metabolic interactions because they offer a direct mechanism for synchronizing growth. Other types of interactions, such as defense or structural support, would require an additional mechanism to link them to reproductive synchronization. Second, we limited our study to endosymbioses between two species to simplify the analyses. Natural endosymbioses can feature potentially multiple partners in different abundances, where synchronizing reproduction would be even more complex and likely depend on system-specific mechanisms. Third, we focused on prokaryote-prokaryote interactions because of the availability and reliability of metabolic models. Although most natural endosymbioses involve eukaryotic hosts and prokaryotic guests, few validated metabolic models exist for eukaryotes. While the analytical results of our mathematical models would still apply, the parameter values drawn from metabolic models would likely differ. In short, the diversity of endosymbioses in nature presents many fascinating opportunities to explore how different forms of interaction influence the evolution of reproductive synchronization.

Here, we focused on reproductive synchronization because it plays an important role in the evolutionary success of endosymbioses. Since synchronization stabilizes the interaction between a host and its guest, it increases the chance that evolution can explore other adaptations, such as various synergies. We show that synchronization may arise relatively easily through simple metabolite sharing and selection for faster growth— it does not require metabolic obligacy or cell cycle coordination. It does, however, require that new endosymbioses are rare. If endosymbioses readily form, then we can paradoxically arrive in a situation where we rarely see host-guest pairs yet they are an important transient stage of the host’s lineage. Moreover, if the formation rate of new endosymbioses drops, we can see the population shift in favor of synchronized endosymbioses. These results highlight that the same mechanism responsible for initiating an endosymbiosis, guest acquisition, is also a barrier to the establishment of synchronization. Thus, the evolutionary stability of endosymbioses may depend on a balance between opportunity and persistence—the faster new partnerships arise, the harder it becomes for any one of them to endure.

## Methods

### Structure of genome-scale metabolic models

We used metabolic models from two of the largest open-access collections available: AGORA (66) and CarveMe (67). As each collection has its own standards for formatting metabolic models, we analyzed the models within each collection separately to prevent errors due to cross-compatibility issues. We used the model files from (44) in.mat format for analysis using MATLAB R2023a (68). In this format, each metabolic model includes a stoichiometric matrix where columns correspond to different metabolic reactions and rows to the different chemical compounds. Reaction fluxes are constrained by lower and upper bounds and a right-hand-side vector indicates how each compound is balanced. All metabolic models come with a default environment so that a metabolism can synthesize all of its biomass compounds (i.e. it is viable). The default environment is determined by the bounds on reaction fluxes and right-hand-side vectors that make a set of extracellular compounds accessible. Determining whether a metabolism is viable requires performing flux balance analysis and solving the associated linear program. We used MATLAB and Gurobi optimization software (69) to solve all linear programs and confirmed that all metabolic models were initially viable, where viability is defined as having an objective function value (growth rate) above a tolerance of 0.001.

To facilitate analyses of endosymbioses, we restructured the metabolic models from AGORA and CarveMe. First, we identified all compounds and reactions used within a collection and reformatted the stoichiometric matrices (*S*) so that the same row corresponds to the same compound across metabolic models. Second, we created an endosymbiotic stoichiometric matrix for each metabolic model depending on its role as either host or guest. A detailed description of how the metabolic models were restructured can be found in (44).

#### Growth rate calculations

We compute the growth rate (fitness) of organisms, either in isolation or endosymbioses, by performing flux balance analysis to solve the associated linear program.

#### Growth rate in isolation

For growth in isolation (free-living), the linear program that calculates the growth rates is:

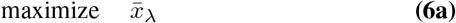

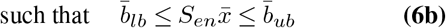

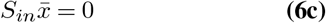

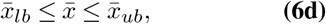

where 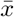 is a vector of reaction fluxes, *λ* is the index for the biomass reaction, and 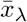 is the flux of the biomass reaction. The first constraint (6b) indicates the exchange of compounds with the environment. The second constraint (6c) indicates that the internal fluxes within cells/organisms are balanced. The third constraint Eq. (6d) indicates the bound on fluxes. *S*_*en*_ and *S*_*in*_ are the stoichiometric matrices for environmental exchanges and intracellular fluxes. The vectors 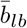 and 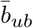 are the lower and upper bounds for derivatives of compound concentrations. The vectors 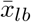 and 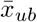 are the lower and upper bounds for fluxes. The outcome of this optimization, 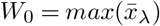, is the growth rate of a free-living organism.

#### Growth rate in a synchronized endosymbiosis

For growth in a synchronized endosymbiosis, the linear program that calculates the growth rates is:

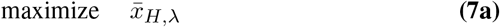

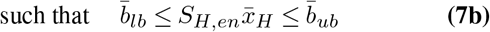

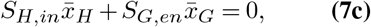

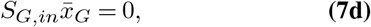

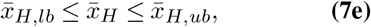

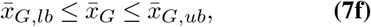

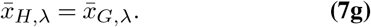

The first constraint in Eq. (7b) is the same as in the isolated growth case. Eq. (7c) indicates that compounds within the host’s intracellular compartments (cytoplasm and periplasm) are balanced. Eq. (7d) indicates that compounds in the guest’s intracellular compartments are balanced. The constraints in Eq. (7e) and Eq. (7f) bound the fluxes of the host and guest. The last constraint Eq. (7g) ensures that the host and guest grow at the same rate. 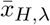 and 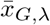 are the host’s and guest’s biomass reactions. *S*_*H,en*_ and *S*_*H,in*_ are the host’s stoichiometric matrices for fluxes with the environment and for intracellular fluxes. *S*_*G,en*_ and *S*_*G,in*_ are the guest’s stoichiometric matrices for fluxes with the environment (which is the host intracellular environment) and for intracellular fluxes. 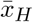 and 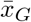 are the host’s and guest’s vectors of reaction fluxes. 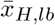 and 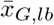 are the host’s and guest’s vectors of lower bounds for fluxes. 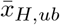 and 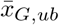 are the host’s and guest’s vectors of upper bounds for fluxes. The outcome of this optimization, 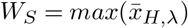, is the synchronized growth rate of a host and its guest.

#### Growth rate in an uncoordinated endosymbiosis

For growth in an uncoordinated endosymbiosis, hosts and guests are allowed to have different growth rates. To calculate these growth rates, we solve the linear program in two steps. In the first step, the linear program that calculates the host’s growth rate is:

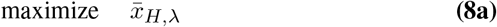

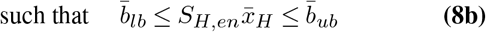

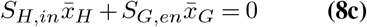

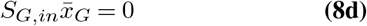

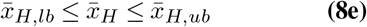

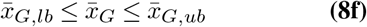

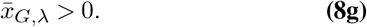

Except for the last constraint Eq. (8g), all other constraints are similar to the case in which there is synchronization. The growth rate of the guest no longer needs to equal that of the host, but it does need to be above zero so as to be a viable endosymbiosis. The outcome of this optimization, 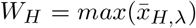, is the growth rate of a host under the uncoordinated growth scenario.

In the second step, the linear program that calculates the guest’s growth rate is:

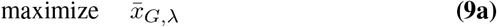

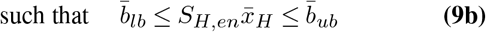

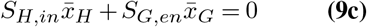

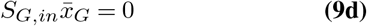

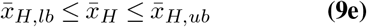

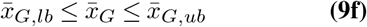

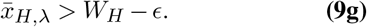

Except for the last constraint Eq. (9g), all other constraints are similar to the case in which there is synchronization. The last constraint requires that the host grow at its maximum as calculated in the first step. To deal with issues of numerical precision we use a small threshold *ϵ* where *ϵ* = 0.001 for the optimization, i.e. *W*_*H*_ − *ϵ*. The outcome of this optimization, 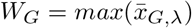, is the growth rate of a guest under the uncoordinated growth scenario.

#### Path to synchronous growth

For a stepwise transition from uncoordinated to synchronous growth, we perform the simulation in four steps (see Fig. 3A). The first and second steps are the same as for growth under the uncoordinated endosymbiosis. These steps will generate the same growth rate as in the uncoordinated scenario. The third and fourth steps are performed iteratively until uncoordinated growth becomes synchronized.

In the third step, the linear program that calculates the host’s growth rate is:

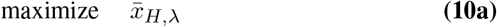

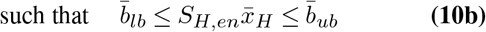

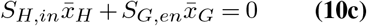

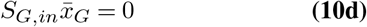

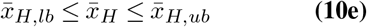

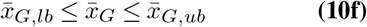

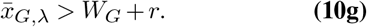

The last constraint requires that the guest’s growth rate be higher than previously calculated by some small amount *r*, set to *r* = 0.001.

In the fourth step, the linear program that calculates the guest’s growth rate is

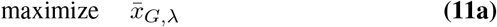

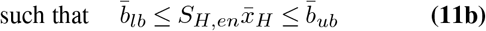

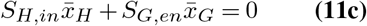

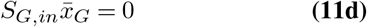

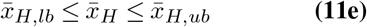

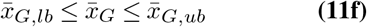

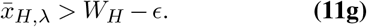

The last step contains the same constraint as the second step, with *W*_*H*_ now representing the values calculated in the third step. We performed the calculations with *ϵ* = 0.001.

## ACKNOWLEDGEMENTS

We are thankful to Kempestiftelserna for the postdoctoral fellowship assigned to Lucas S. Souza (SMK21-0004).

## Supporting information

**Fig. S1.**
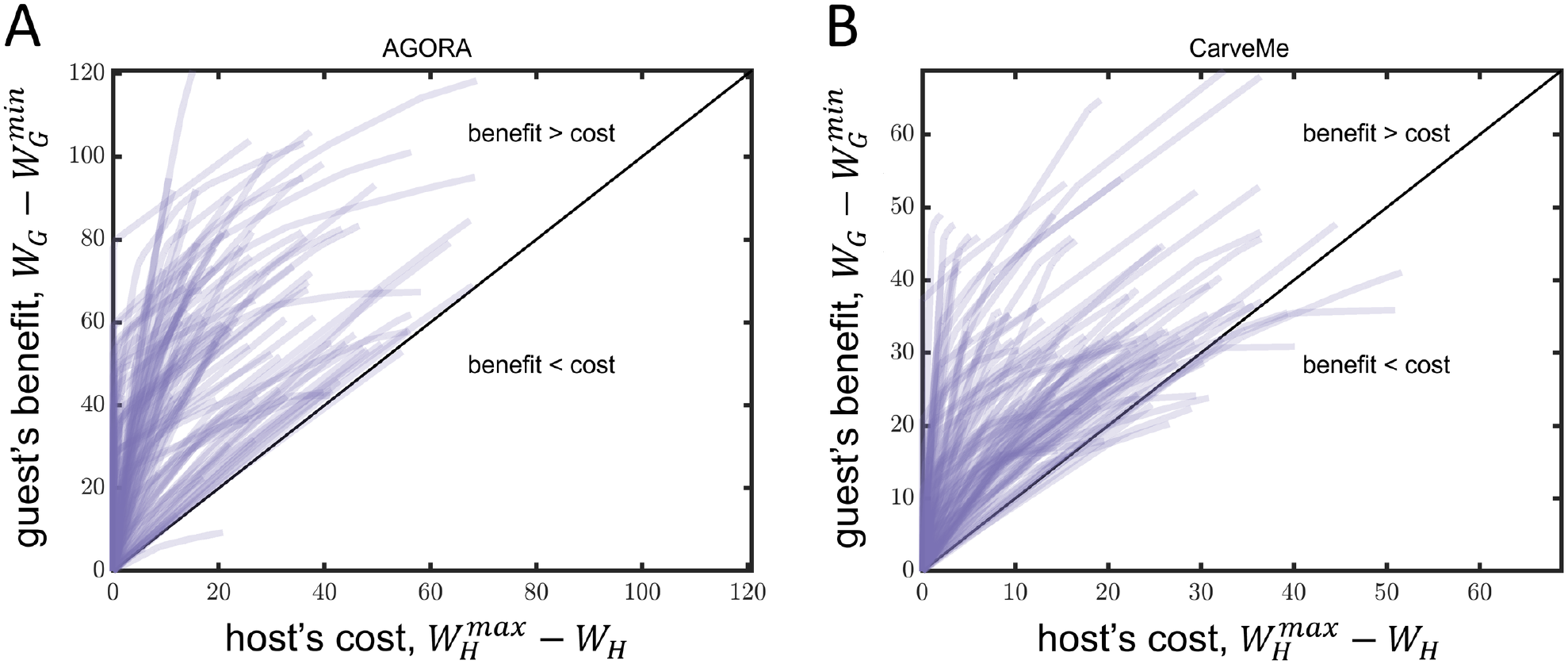
Trade-off between guest’s benefit and host’s cost under increasing synchronization. (A-B) Each graph shows 100 curves, each corresponding to the simulation of a distinct pair (the same pairs from Figs. 3B-D) randomly selected from those where *W*_*H*_ > *W*_*S*_ > *W*_*G*_. Pairs from AGORA (A) and CarveMe (B) are distinct. The leftmost values of hosts’ and guests’ fitness are under the completely uncoordinated scenario, while the rightmost values correspond to the fully synchronized scenario. Intermediate values represent progressive synchronization of hosts’ and guests’ fitness. The host’s fitness is the highest 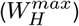, and the guest’s fitness is the lowest 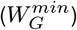 at the leftmost values. We name ‘guest’s benefit’ the gained growth of guests from a stage of the uncoordinated scenario to an increase in synchronization 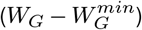. Similarly, we name ‘host’s cost’ the lost growth of hosts from a stage of the uncoordinated scenario to an increase in synchronization 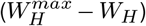. Both graphs show an asymmetry in fitnesses, with the majority of cases having the guest’s benefit surpassing the host’s cost within the whole range of synchronization.

**Fig. S2.**
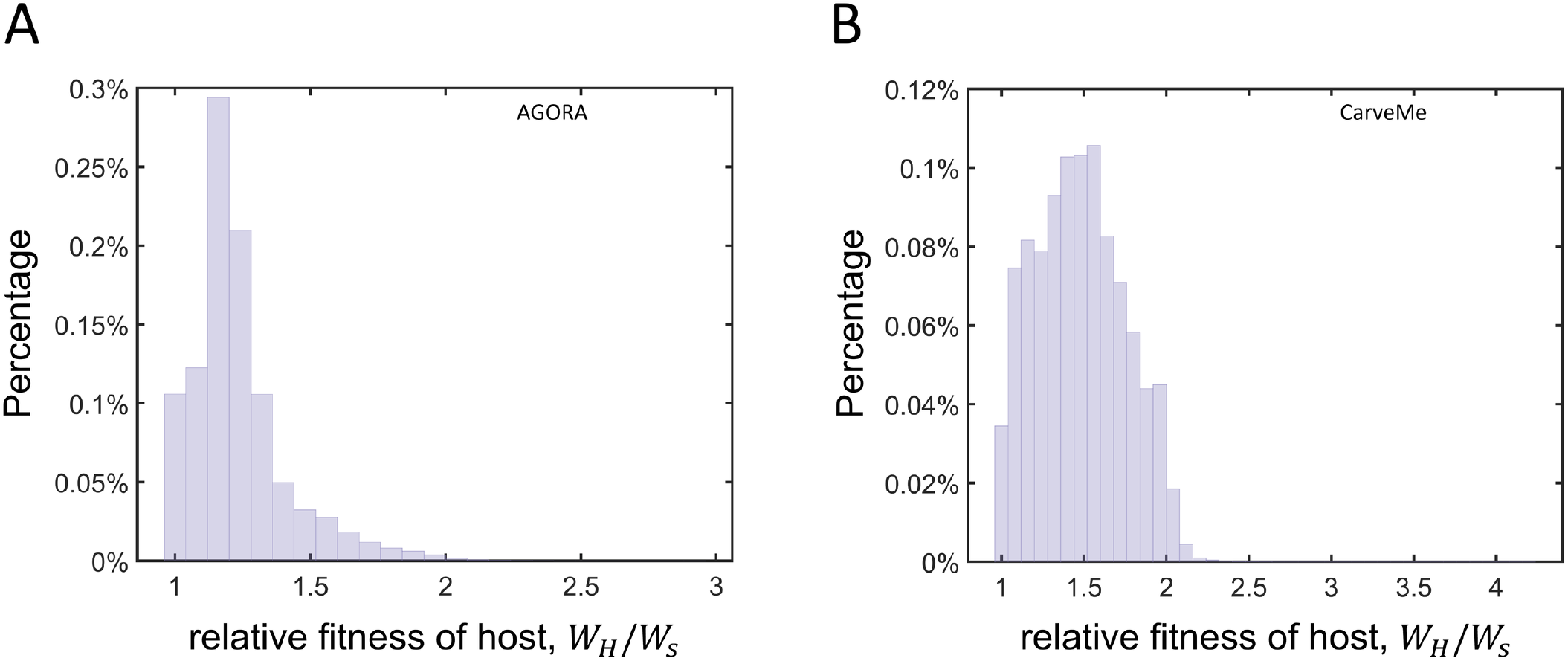
Histograms of relative fitness advantage of uncoordinated hosts versus synchronized hosts. (A-B) The histograms show the distribution of uncoordinated host fitness (*W*_*H*_) relative to synchronized host fitness (*W*_*S*_), estimated from flux balance analysis. (A) For AGORA, uncoordinated hosts grow on average 19% faster than synchronized hosts, median(*W*_*H*_ /*W*_*S*_) = 1.1936, and this advantage spans from 1.001 to 2.9354. The 75th, 90th, and 99th percentiles are 1.2894, 1.4655, and 1.8804, respectively. (B) For CarveMe, uncoordinated hosts grow on average 47% faster than synchronized hosts, median(*W*_*H*_ /*W*_*S*_) = 1.4681, and this advantage spans from 1.001 to 4.1865. The 75th, 90th, and 99th percentiles are 1.6731, 1.8634, and 2.0576, respectively. (A-B) Both histograms show that uncoordinated hosts have higher fitness than synchronizing hosts. The histograms for AGORA and CarveMe have 53, 591 and 2, 452, 637 distinct host-guest pairs, respectively. We used all pairs for which synchronized hosts outcompete both free-living and uncoordinated hosts, accounting for the dissociation rate and assuming the formation of new endosymbioses is rare.

**Fig. S3.**
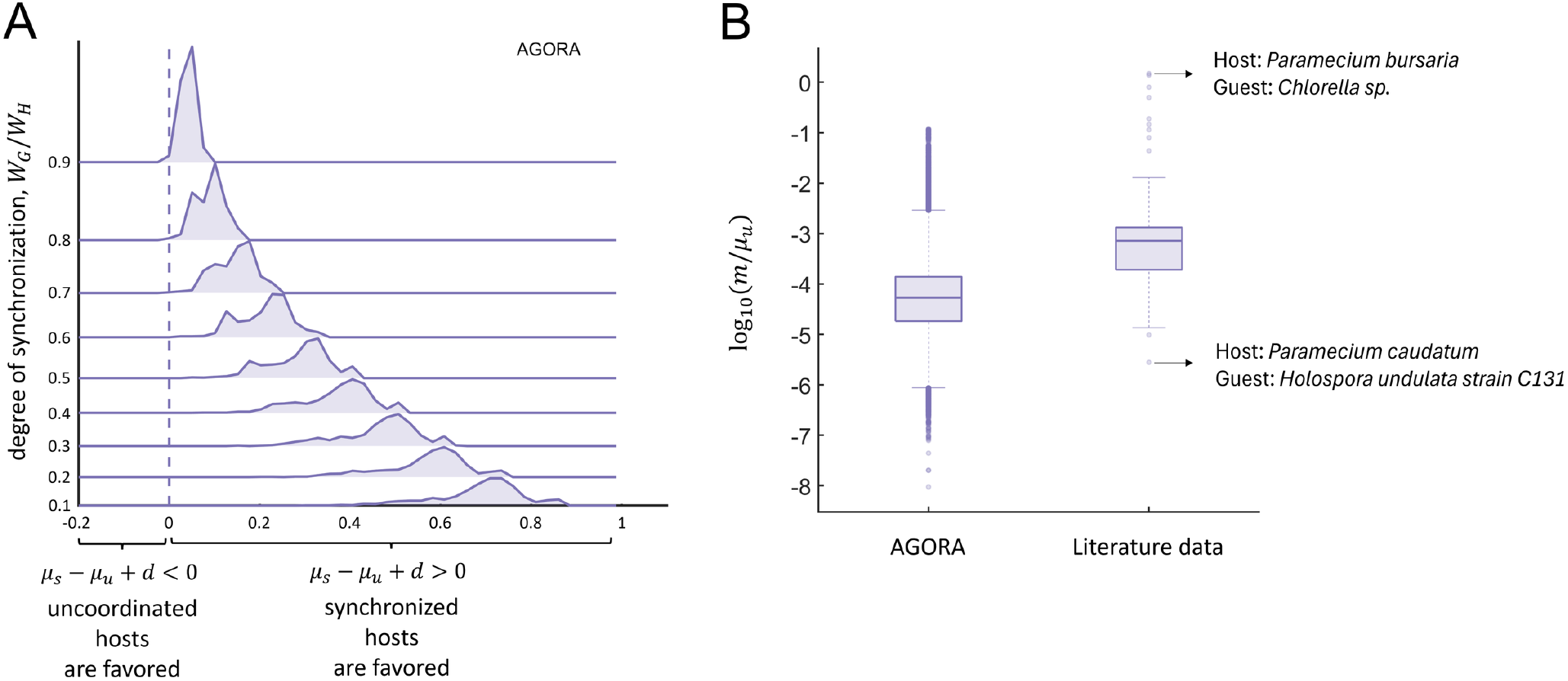
Companion to Fig. 4 using AGORA models. (A) A joyplot displays a series of histograms for different degrees of synchronization (*W*_*G*_/*W*_*H*_). Each histogram shows the distribution of the differences in net growth rate between synchronized (*µ*_*s*_) and uncoordinated growth (*µ*_*u*_ −*d*) in 1000 host-guest pairs using AGORA models (the corresponding graph for CarveMe is Fig. 4B). The dashed line separates where uncoordinated growth (left) and synchronization (right) are favored. In the vast majority of host-guest pairs studied, synchronization results in higher growth. (B) Boxplots show the distribution of the endosymbioses formation rate (*m*) scaled by uncoordinated host growth rate from two different datasets: 1. our modeling analyses (labeled AGORA) and 2. data in the literature. For the AGORA boxplot, we used the critical rate at which synchronization is disfavored *m*^*\**^ (Eq. (5)) as the value of *m*, with *ϕ* = 0.01. For the literature data boxplot, we calculated *m* as described in *SI-B* and we used *µ*_*u*_ = *ln*2/24, with 24 denoting the doubling time (in hours) of the population. In each box, the central mark indicates the median, the bottom and top edges represent the 25th and 75th percentiles, and the whiskers extend to the most extreme data points not considered outliers, which are plotted as individual dots. The AGORA boxplot uses 53,591 measurements (all distinct host–guest pairs where synchronizing hosts are fitter when *m* = 0) and the literature data boxplot uses 78 measurements. The corresponding graph for CarveMe is found in Fig. 4D. Values from the experimental observations are higher on average than those from our modeling analyses.

**Fig. S4.**
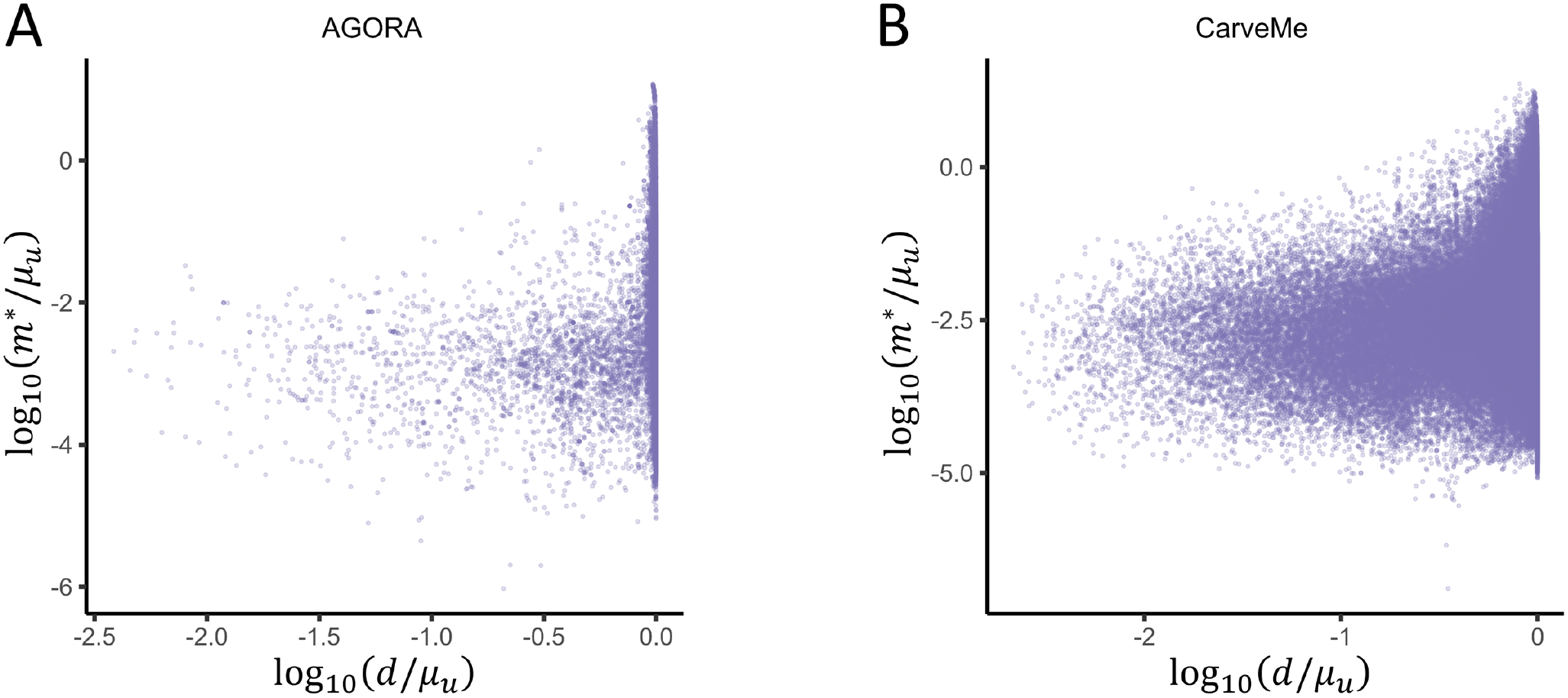
Association between the critical formation rate of endosymbiosis and dissociation rate. (A-B) The scatter plots show the critical formation rate of endosymbiosis (*m*\*) versus dissociation rate (*d*)—both normalized by the growth rate of uncoordinated hosts (*µ*_*u*_)— on a logarithmic scale. The measurements from AGORA and CarveMe are the same as those in Fig. S3B and Fig. 4D, respectively. The AGORA and CarveMe scatter plots use 53,591 and 2,452,637 measurements, respectively, each representing distinct host–guest pairs where synchronizing hosts are fitter when the formation of new endosymbiosis is sufficiently rare. Both scatter plots show only a weak association between the dissociation and formation of new endosymbiosis.

**Table S1.** Joint distribution of viability under synchronized and uncoordinated endosymbiosis for AGORA.

|  |  | **Uncoordinated** |  |
| --- | --- | --- | --- |
|  |  | Viable | Non-Viable |
| **Synchronized** | Viable | 0.5561 | 0.0030 |
|  | Non-Viable | 0.0070 | 0.4339 |

**Table S2.** Joint distribution of viability under synchronized and uncoordinated endosymbiosis for CarveMe.

|  |  | **Uncoordinated** |  |
| --- | --- | --- | --- |
|  |  | Viable | Non-Viable |
| **Synchronized** | Viable | 0.5814 | 0.0396 |
|  | Non-Viable | 0.0021 | 0.3769 |

**Table S3.** List of variables and parameters associated with our system of differential equations.

| Symbol | Description | Range/constraints | Unit |
| --- | --- | --- | --- |
| $R$ | Resource concentration available to all hosts | $R \geq 0$ | concentration of resource: [R] |
| $H_u, H_s, H_0$ | Population densities of uncoordinated hosts ($H_u$), synchronized hosts ($H_s$), and host without guest ($H_0$) | $H_u, H_s, H_0 \geq 0$ | density (e.g., cells · volume<sup>-1</sup>) |
| $\mu_u, \mu_s, \mu_0$ | Maximum growth rates for $H_u$, $H_s$, and $H_0$ | $\mu_u > \mu_s > \mu_0 > 0$ | time<sup>-1</sup> |
| $d$ | Dissociation rate ($H_u \rightarrow H_0$) | $d \geq 0$ | time<sup>-1</sup> |
| $m$ | Formation rate of new endosymbioses ($H_0 \rightarrow H_u$) | $m \geq 0$ | time<sup>-1</sup> |
| $c$ | Per-host resource consumption rate | $c > 0$ | [R]/(host · time) |
| $k_r$ | Inflow/supply rate of resource into the environment | $k_r > 0$ | [R]/time |
| $\theta$ | Monod half-saturation constant controlling resource limitation | $\theta > 0$ | [R] |
| $\phi$ | Outflux rate affecting resource and all hosts | $\phi > 0$ | time<sup>-1</sup> |

**Table S4.**
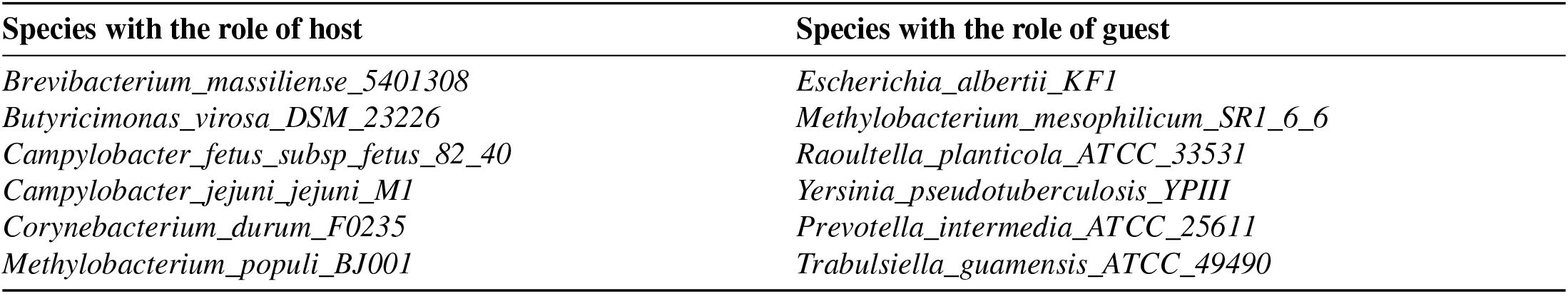
This table lists species within AGORA that form host–guest pairs in which growth rates under the uncoordinated scenario naturally led to synchronization. In total, 6 distinct pairs were identified out of all possible pair combinations that are viable (372,120 pairs), corresponding to the ‘< 0.01%’ shown in Fig. 2B.

**Table S5.**
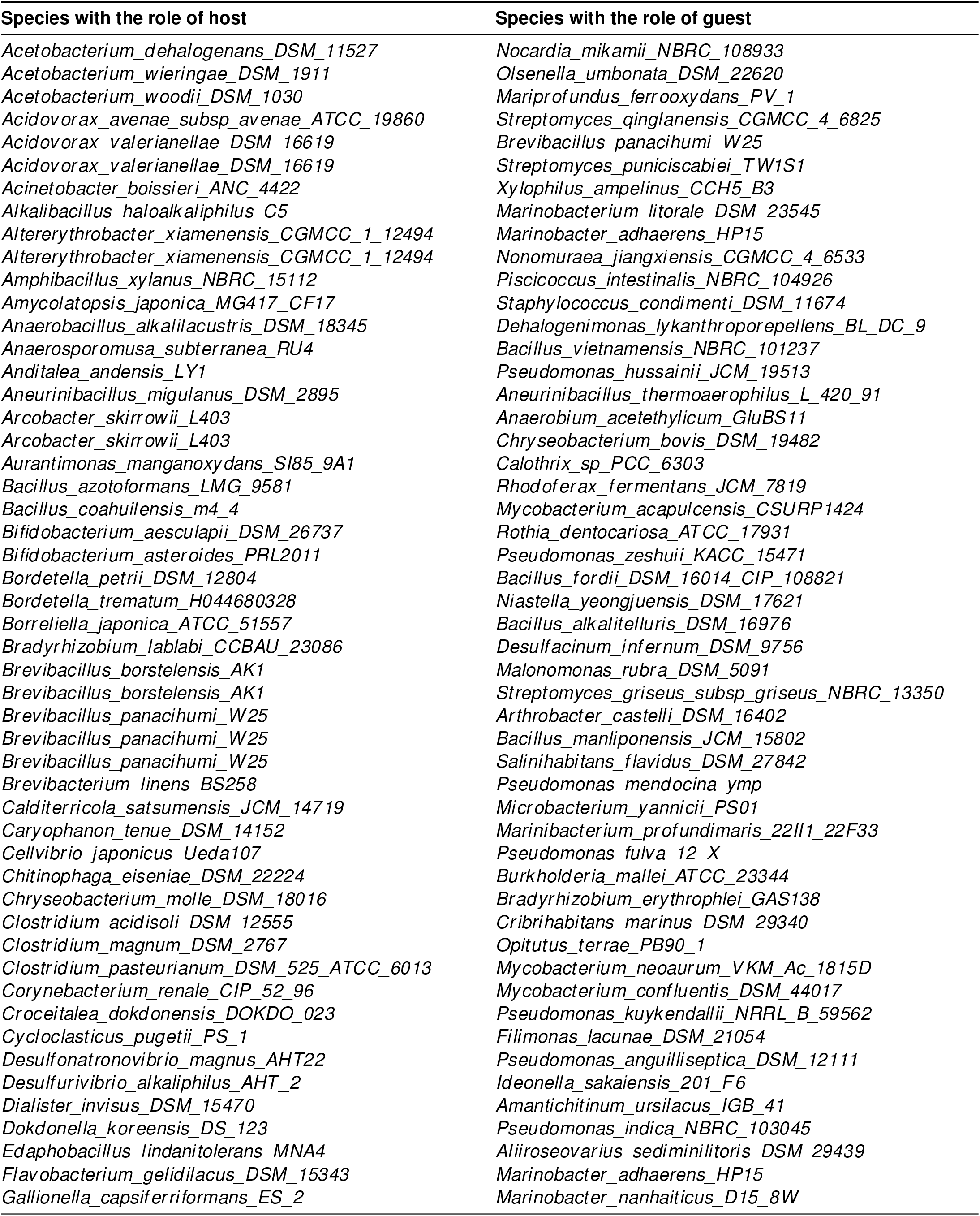
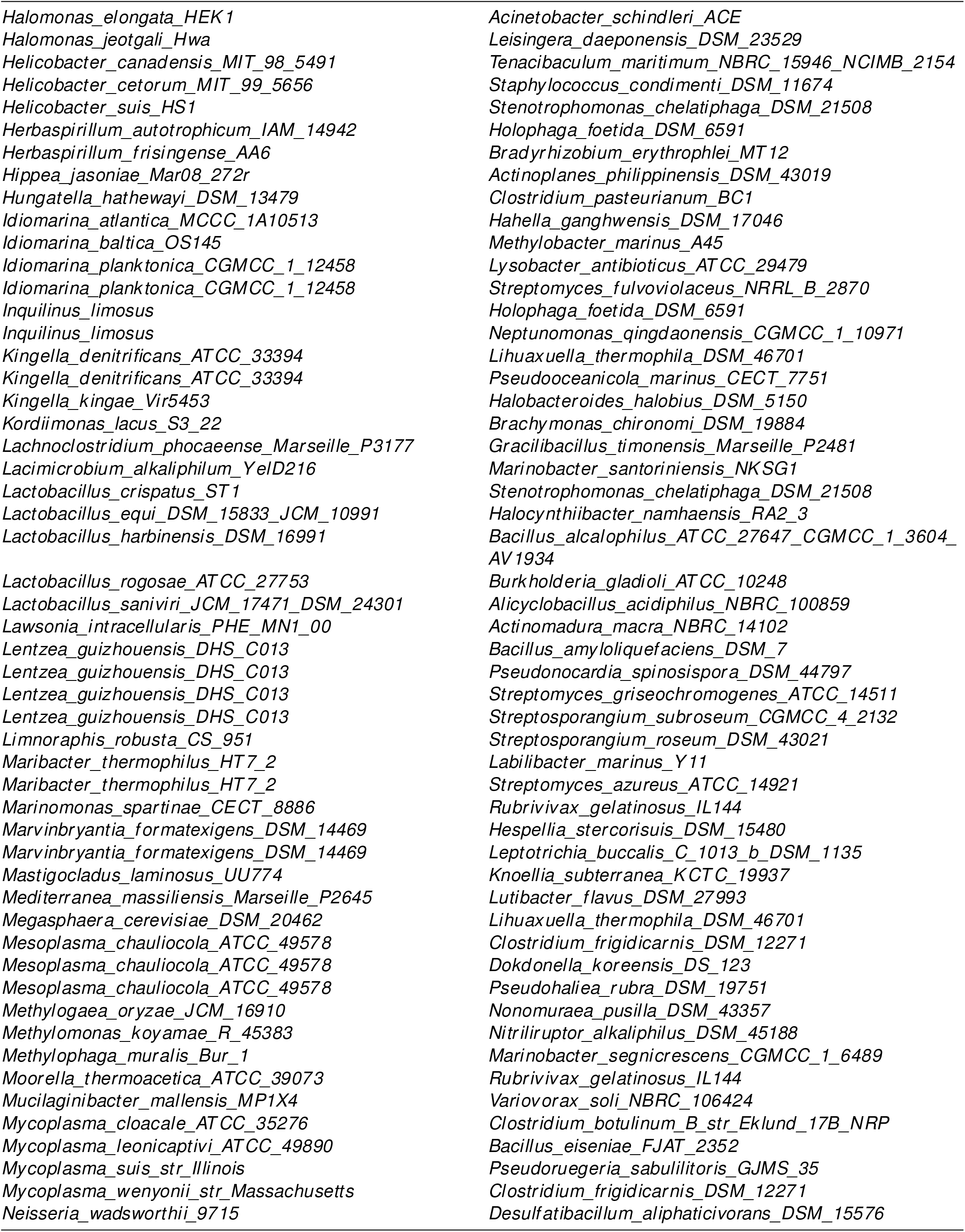
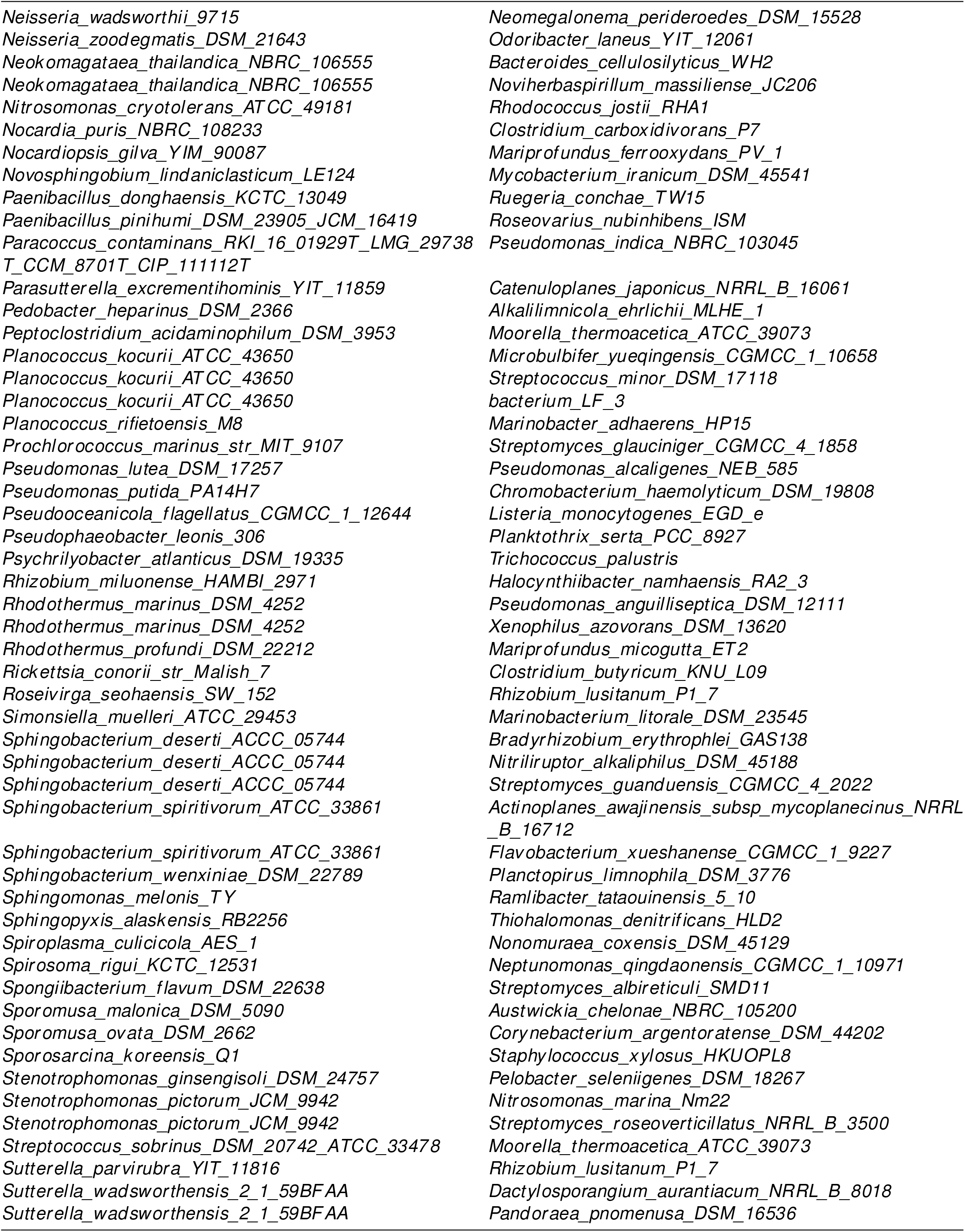
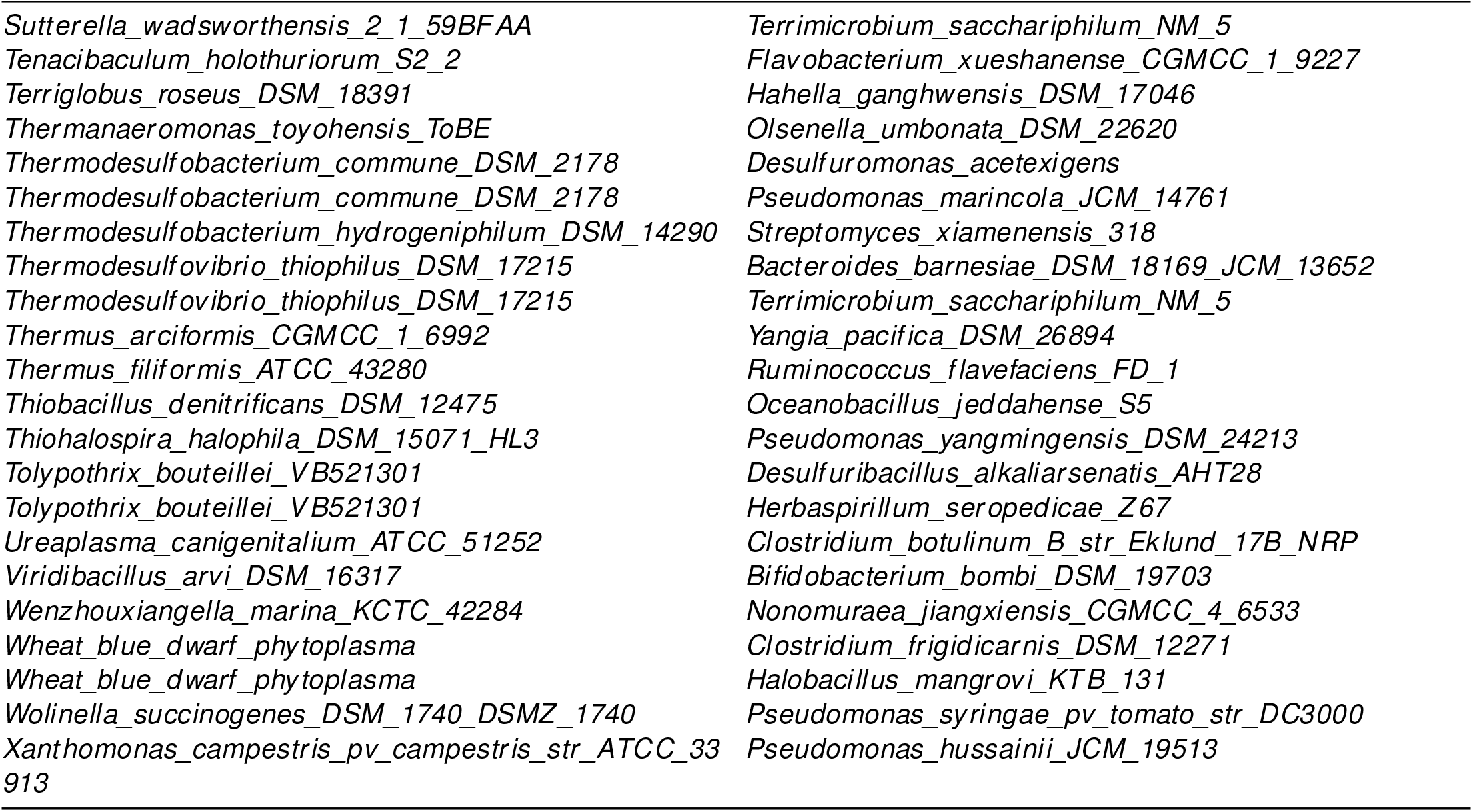
This table lists species within CarveMe that form host–guest pairs in which growth rates under the uncoordinated scenario naturally led to synchronization. In total, 178 distinct pairs were identified out of all possible pair combinations that are viable (18,147,674 pairs), corresponding to ‘< 0.001%’ as shown in Fig. 2B.

**Table S6.**
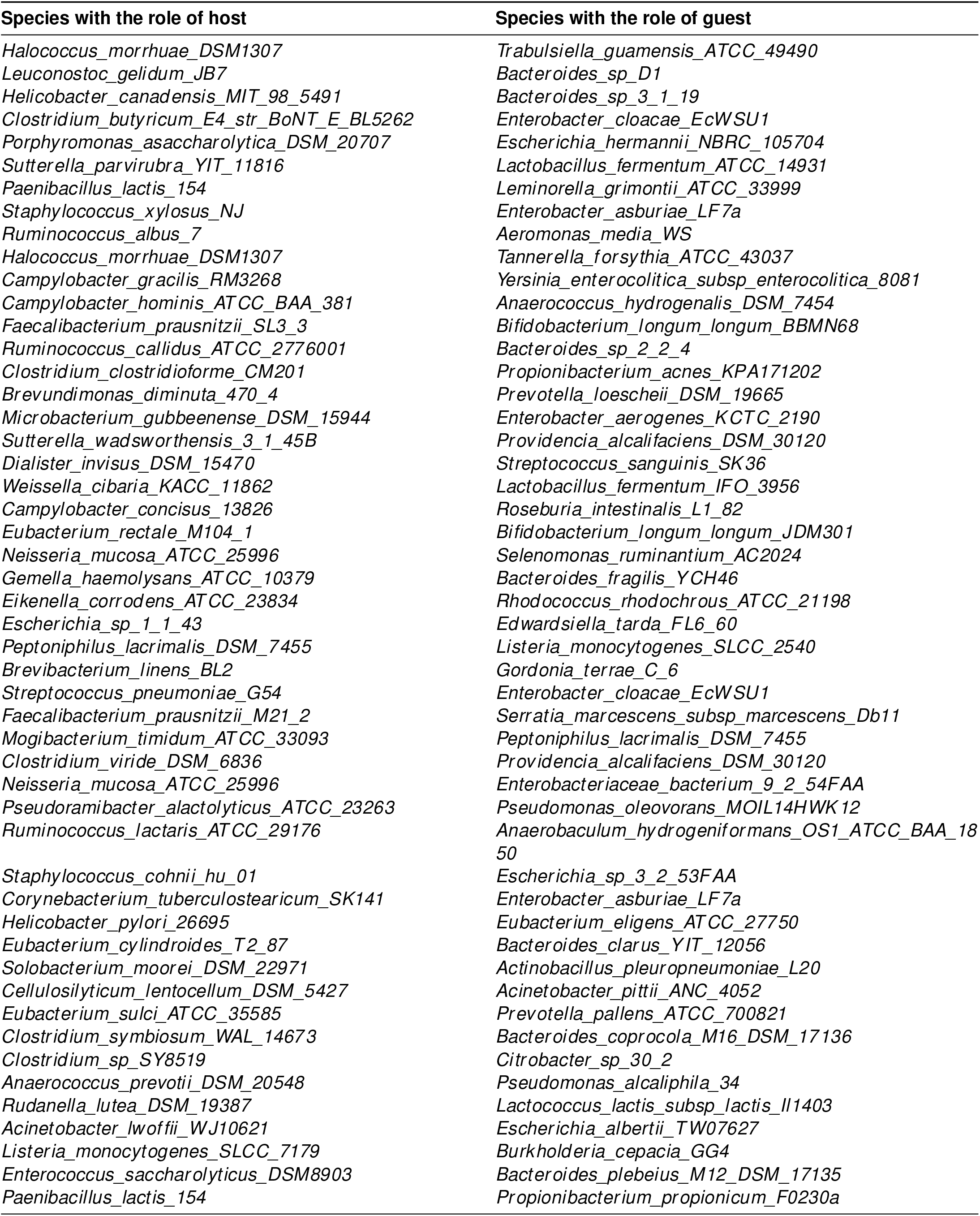
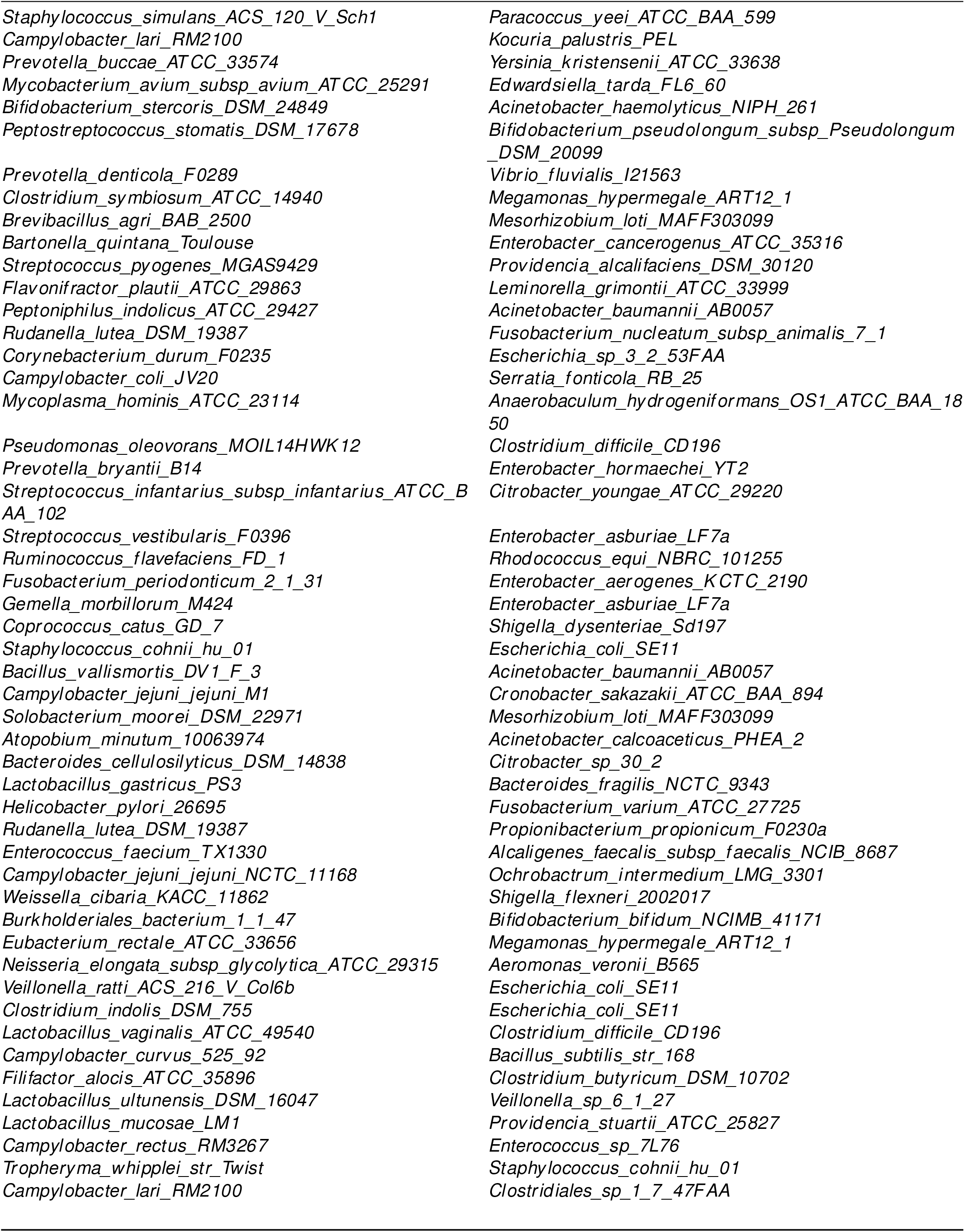
This table lists the 100 distinct host–guest pairs from AGORA used to illustrate how an initially uncoordinated pair can become synchronized by iteratively requiring hosts to increase their guests’ growth. Data from these pairs were used to generate Fig. 3B–D.

| Species with the role of host | Species with the role of guest |
| --- | --- |
| *Halococcus_morruhae_DSM1307* | *Trabulsiella_guamensis_ATCC_49490* |
| *Leuconostoc_gelidum_JB7* | *Bacteroides_sp_D1* |
| *Helicobacter_canadensis_MIT_98_5491* | *Bacteroides_sp_3_1_19* |
| *Clostridium_butyricum_E4_str_BoNT_E_BL5262* | *Enterobacter_cloacae_EcWSU1* |
| *Porphyromonas_asaccharolytica_DSM_20707* | *Escherichia_hermannii_NBRC_105704* |
| *Sutterella_parvirubra_YIT_11816* | *Lactobacillus_fermentum_ATCC_14931* |
| *Paenibacillus_lactis_154* | *Leminorella_grimontii_ATCC_33999* |
| *Staphylococcus_xylosum_NJ* | *Enterobacter_asburiae_LF7a* |
| *Ruminococcus_albus_7* | *Aeromonas_media_WS* |
| *Halococcus_morruhae_DSM1307* | *Tannerella_forsythia_ATCC_43037* |
| *Campylobacter_gracilis_RM3268* | *Yersinia_enterocolitica_subsp_enterocolitica_8081* |
| *Campylobacter_hominis_ATCC_BAA_381* | *Anaerococcus_hydrogenalis_DSM_7454* |
| *Faecalibacterium_prausnitzii_SL3_3* | *Bifidobacterium_longum_longum_BBMN68* |
| *Ruminococcus_callidus_ATCC_2776001* | *Bacteroides_sp_2_2_4* |
| *Clostridium_clostridioforme_CM201* | *Propionibacterium_acnes_KPA171202* |
| *Brevundimonas_diminuta_470_4* | *Prevotella_loeschei_DSM_19665* |
| *Microbacterium_gubbeenense_DSM_15944* | *Enterobacter_aerogenes_KCTC_2190* |
| *Sutterella_wadsworthensis_3_1_45B* | *Providencia_alcalifaciens_DSM_30120* |
| *Dialister_invisus_DSM_15470* | *Streptococcus_sanguinis_SK36* |
| *Weissella_cibaria_KACC_11862* | *Lactobacillus_fermentum_IFO_3956* |
| *Campylobacter_conciscus_13826* | *Roseburia_intestinalis_L1_82* |
| *Eubacterium_rectale_M104_1* | *Bifidobacterium_longum_longum_JDM301* |
| *Neisseria_mucosa_ATCC_25996* | *Selenomonas_ruminantium_AC2024* |
| *Gemella_haemolysans_ATCC_10379* | *Bacteroides_fragilis_YCH46* |
| *Eikenella_corrodens_ATCC_23834* | *Rhodococcus_rhodochrous_ATCC_21198* |
| *Escherichia_sp_1_1_43* | *Edwardsiella_tarda_FL6_60* |
| *Peptoniphilus_lacrimalis_DSM_7455* | *Listeria_monocytogenes_SLCC_2540* |
| *Brevibacterium_linens_BL2* | *Gordonia_terrae_C_6* |
| *Streptococcus_pneumoniae_G54* | *Enterobacter_cloacae_EcWSU1* |
| *Faecalibacterium_prausnitzii_M21_2* | *Serratia_marcescens_subsp_marcescens_Db11* |
| *Mogibacterium_timidum_ATCC_33093* | *Peptoniphilus_lacrimalis_DSM_7455* |
| *Clostridium_viride_DSM_6836* | *Providencia_alcalifaciens_DSM_30120* |
| *Neisseria_mucosa_ATCC_25996* | *Enterobacteriaceae_bacterium_9_2_54FAA* |
| *Pseudoramibacter_alactolyticus_ATCC_23263* | *Pseudomonas_oleovorans_MOIL14HWK12* |
| *Ruminococcus_lactaris_ATCC_29176* | *Anaerobaculum_hydrogeniformans_OS1_ATCC_BAA_18_50* |
| *Staphylococcus_cohnii_hu_01* | *Escherichia_sp_3_2_53FAA* |
| *Corynebacterium_tuberculoostearicum_SK141* | *Enterobacter_asburiae_LF7a* |
| *Helicobacter_pylori_26695* | *Eubacterium_eligens_ATCC_27750* |
| *Eubacterium_cylindroides_T2_87* | *Bacteroides_clarus_YIT_12056* |
| *Solobacterium_moorei_DSM_22971* | *Actinobacillus_pleuropneumoniae_L20* |
| *Cellulosilyticum_lentocellum_DSM_5427* | *Acinetobacter_pittii_ANC_4052* |
| *Eubacterium_sulci_ATCC_35585* | *Prevotella_pallens_ATCC_700821* |
| *Clostridium_symbiosum_WAL_14673* | *Bacteroides_coprocola_M16_DSM_17136* |
| *Clostridium_sp_SY8519* | *Citrobacter_sp_30_2* |
| *Anaerococcus_prevotii_DSM_20548* | *Pseudomonas_alcaliphila_34* |
| *Rudanella_lutea_DSM_19387* | *Lactococcus_lactis_subsp_lactis_II1403* |
| *Acinetobacter_lwoffii_WJ10621* | *Escherichia_albertii_TW07627* |
| *Listeria_monocytogenes_SLCC_7179* | *Burkholderia_cepacia_GG4* |
| *Enterococcus_saccharolyticus_DSM8903* | *Bacteroides_plebeius_M12_DSM_17135* |
| *Paenibacillus_lactis_154* | *Propionibacterium_propionicum_F0230a* |
| *Staphylococcus_simulans_ACS_120_V_Sch1* | *Paracoccus_yeei_ATCC_BAA_599* |
| *Campylobacter_lari_RM2100* | *Kocuria_palustris_PEL* |
| *Prevotella_buccae_ATCC_33574* | *Yersinia_kristensenii_ATCC_33638* |
| *Mycobacterium_avium_subsp_avium_ATCC_25291* | *Edwardsiella_tarda_FL6_60* |
| *Bifidobacterium_stercoris_DSM_24849* | *Acinetobacter_haemolyticus_NIPH_261* |
| *Peptostreptococcus_stomatis_DSM_17678* | *Bifidobacterium_pseudolongum_subsp_Pseudolongum_DSM_20099* |
| *Prevotella_denticola_F0289* | *Vibrio_fluvialis_I21563* |
| *Clostridium_symbiosum_ATCC_14940* | *Megamonas_hypermegale_ART12_1* |
| *Brevibacillus_agri_BAB_2500* | *Mesorhizobium_loti_MAFF303099* |
| *Bartonella_quintana_Toulouse* | *Enterobacter_cancerogenus_ATCC_35316* |
| *Streptococcus_pyogenes_MGAS9429* | *Providencia_alcalifaciens_DSM_30120* |
| *Flavonifractor_plautii_ATCC_29863* | *Leminorella_grimontii_ATCC_33999* |
| *Peptoniphilus_indolicus_ATCC_29427* | *Acinetobacter_baumannii_AB0057* |
| *Rudanella_lutea_DSM_19387* | *Fusobacterium_nucleatum_subsp_animalis_7_1* |
| *Corynebacterium_durum_F0235* | *Escherichia_sp_3_2_53FAA* |
| *Campylobacter_coli_JV20* | *Serratia_fonticola_RB_25* |
| *Mycoplasma_hominis_ATCC_23114* | *Anaerobaculum_hydrogeniformans_OS1_ATCC_BAA_18_50* |
| *Pseudomonas_oleovorans_MOIL14HWK12* | *Clostridium_difficile_CD196* |
| *Prevotella_bryantii_B14* | *Enterobacter_hormaechei_YT2* |
| *Streptococcus_infantarius_subsp_infantarius_ATCC_BAA_102* | *Citrobacter_youngae_ATCC_29220* |
| *Streptococcus_vestibularis_F0396* | *Enterobacter_asburiae_LF7a* |
| *Ruminococcus_flavofaciens_FD_1* | *Rhodococcus_equi_NBRC_101255* |
| *Fusobacterium_periodonticum_2_1_31* | *Enterobacter_aerogenes_KCTC_2190* |
| *Gemella_morbillorum_M424* | *Enterobacter_asburiae_LF7a* |
| *Coprococcus_catus_GD_7* | *Shigella_dysenteriae_Sd197* |
| *Staphylococcus_cohnii_hu_01* | *Escherichia_coli_SE11* |
| *Bacillus_vallismortis_DV1_F_3* | *Acinetobacter_baumannii_AB0057* |
| *Campylobacter_jejuni_jejuni_M1* | *Cronobacter_sakazakii_ATCC_BAA_894* |
| *Solobacterium_moorei_DSM_22971* | *Mesorhizobium_loti_MAFF303099* |
| *Atopobium_minutum_10063974* | *Acinetobacter_calcoaceticus_PHEA_2* |
| *Bacteroides_cellulosilyticus_DSM_14838* | *Citrobacter_sp_30_2* |
| *Lactobacillus_gastricus_PS3* | *Bacteroides_fragilis_NCTC_9343* |
| *Helicobacter_pylori_26695* | *Fusobacterium_varium_ATCC_27725* |
| *Rudanella_lutea_DSM_19387* | *Propionibacterium_propionicum_F0230a* |
| *Enterococcus_faecium_TX1330* | *Alcaligenes_faecalis_subsp_faecalis_NCIB_8687* |
| *Campylobacter_jejuni_jejuni_NCTC_11168* | *Ochrobactrum_intermedium_LMG_3301* |
| *Weissella_cibaria_KACC_11862* | *Shigella_flexneri_2002017* |
| *Burkholderiales_bacterium_1_1_47* | *Bifidobacterium_bifidum_NCIMB_41171* |
| *Eubacterium_rectale_ATCC_33656* | *Megamonas_hypermegale_ART12_1* |
| *Neisseria_elongata_subsp_glycolytica_ATCC_29315* | *Aeromonas_veronii_B565* |
| *Veillonella_ratti_ACS_216_V_Col6b* | *Escherichia_coli_SE11* |
| *Clostridium_indolis_DSM_755* | *Escherichia_coli_SE11* |
| *Lactobacillus_vaginalis_ATCC_49540* | *Clostridium_difficile_CD196* |
| *Campylobacter_curvus_525_92* | *Bacillus_subtilis_str_168* |
| *Filifactor_alocis_ATCC_35896* | *Clostridium_butyricum_DSM_10702* |
| *Lactobacillus_ultunensis_DSM_16047* | *Veillonella_sp_6_1_27* |
| *Lactobacillus_mucosae_LM1* | *Providencia_stuartii_ATCC_25827* |
| *Campylobacter_rectus_RM3267* | *Enterococcus_sp_7L76* |
| *Tropheryma_whipplei_str_Twist* | *Staphylococcus_cohnii_hu_01* |
| *Campylobacter_lari_RM2100* | *Clostridiales_sp_1_7_47FAA* |

**Table S7.**
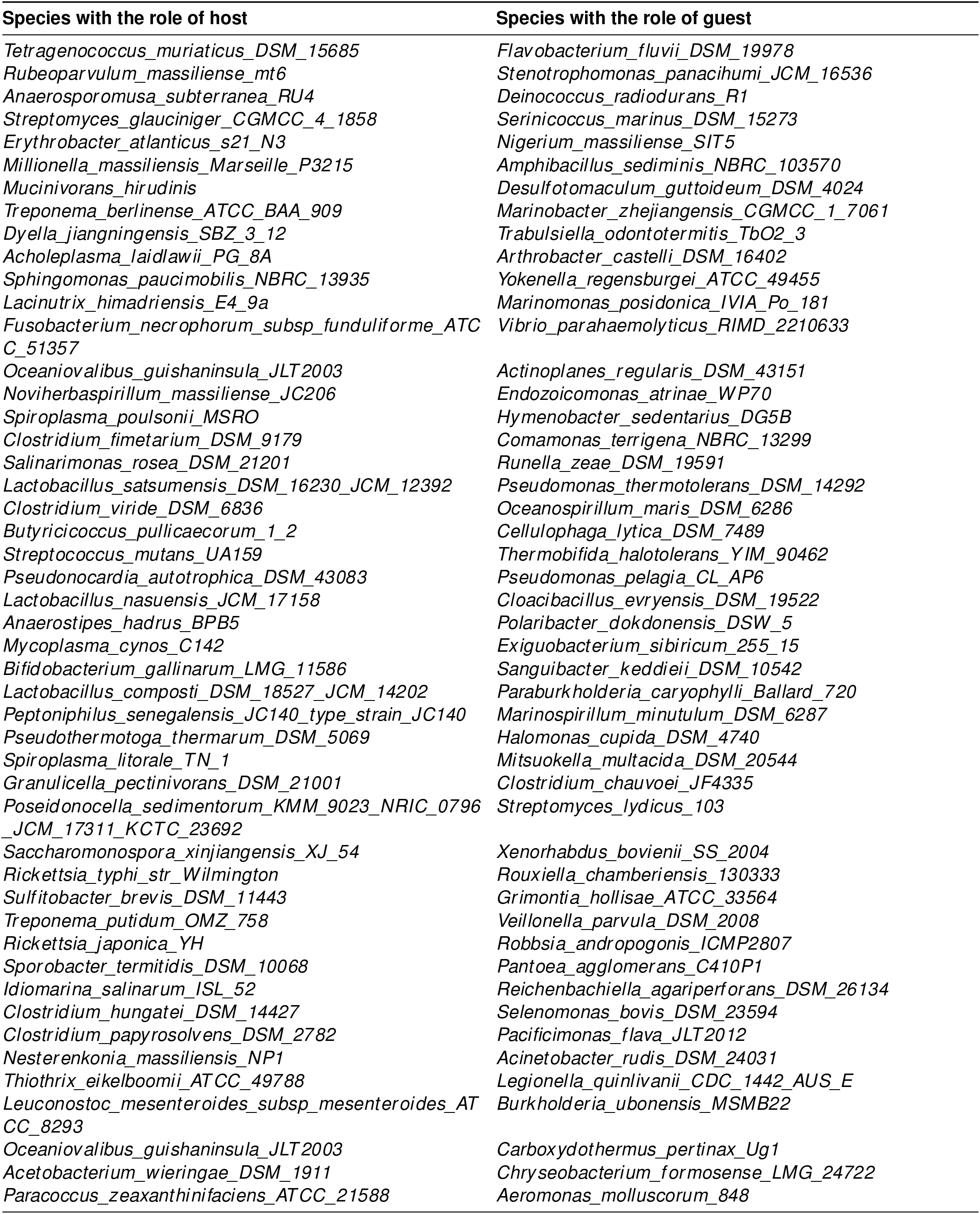
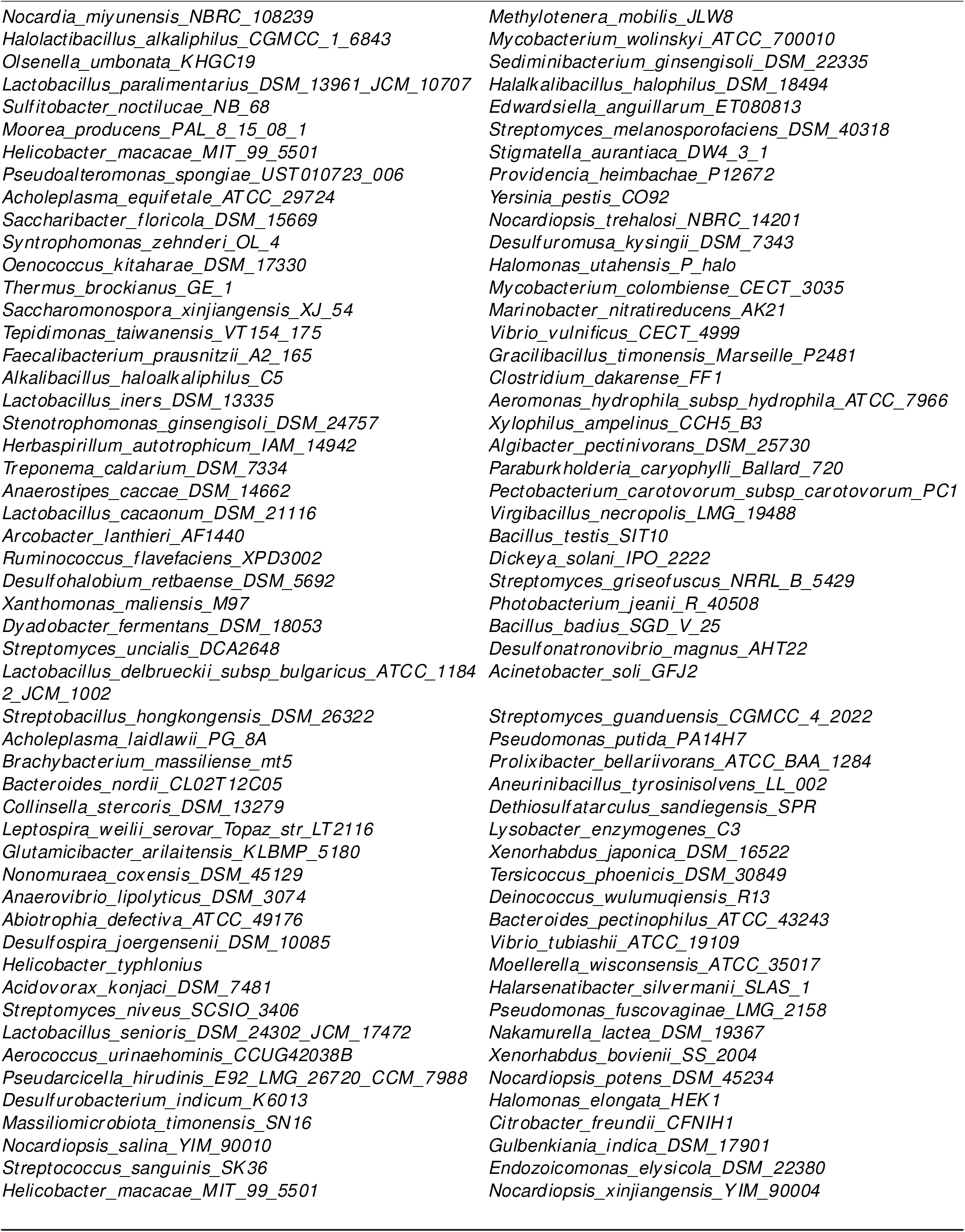
This table lists the 100 distinct host–guest pairs from CarveMe used to illustrate how an initially uncoordinated pair can become synchronized by iteratively requiring hosts to increase their guests’ growth. Data from these pairs were used to generate Fig. 3B–D.

| Species with the role of host | Species with the role of guest |
| --- | --- |
| *Tetragenococcus muriaticus*_DSM_15685 | *Flavobacterium fluvii*_DSM_19978 |
| *Rubeoparvulum massiliense*_mt6 | *Stenotrophomonas panacihumi*_JCM_16536 |
| *Anaerosporemusa subterranea*_RU4 | *Deinococcus radiodurans*_R1 |
| *Streptomyces glauciniger*_CGMCC_4_1858 | *Serinicoccus marinus*_DSM_15273 |
| *Erythrobacter atlanticus*_s21_N3 | *Nigerium massiliense*_SIT5 |
| *Millionella massiliensis*_Marseille_P3215 | *Amphibacillus sediminis*_NBRC_103570 |
| *Mucinivorans hirudinis* | *Desulfotomaculum guttoideum*_DSM_4024 |
| *Treponema berlinense*_ATCC_BAA_909 | *Marinobacter zhejiangensis*_CGMCC_1_7061 |
| *Dyella jiangningensis*_SBZ_3_12 | *Trabulsiella odontotermitis*_TbO2_3 |
| *Acholeplasma laidlawii*_PG_8A | *Arthrobacter castelli*_DSM_16402 |
| *Sphingomonas paucimobilis*_NBRC_13935 | *Yokenella regensburgei*_ATCC_49455 |
| *Lacinutrix himadriensis*_E4_9a | *Marinomonas posidonica*_IVIA_Po_181 |
| *Fusobacterium necrophorum*_subsp_funduliforme_ATC C_51357 | *Vibrio parahaemolyticus*_RIMD_2210633 |
| *Oceaniovalibus guishaninsula*_JLT2003 | *Actinoplanes regularis*_DSM_43151 |
| *Noviherbaspirillum massiliense*_JC206 | *Endozoicomonas atrinae*_WP70 |
| *Spiroplasma poulsonii*_MSRO | *Hymenobacter sedentarius*_DG5B |
| *Clostridium fimetarium*_DSM_9179 | *Comamonas terrigena*_NBRC_13299 |
| *Salinarimonas rosea*_DSM_21201 | *Runella zeae*_DSM_19591 |
| *Lactobacillus satsumensis*_DSM_16230_JCM_12392 | *Pseudomonas thermotolerans*_DSM_14292 |
| *Clostridium viride*_DSM_6836 | *Oceanospirillum maris*_DSM_6286 |
| *Butyricicoccus pullicaecorum*_1_2 | *Cellulophaga lytica*_DSM_7489 |
| *Streptococcus mutans*_UA159 | *Thermobifida halotolerans*_YIM_90462 |
| *Pseudonocardia autotrophica*_DSM_43083 | *Pseudomonas pelagia*_CL_AP6 |
| *Lactobacillus nasuensis*_JCM_17158 | *Cloacibacillus evryensis*_DSM_19522 |
| *Anaerostipes hadrus*_BPB5 | *Polaribacter dokdonensis*_DSW_5 |
| *Mycoplasma cynos*_C142 | *Exiguobacterium sibiricum*_255_15 |
| *Bifidobacterium gallinarum*_LMG_11586 | *Sanguibacter keddiei*_DSM_10542 |
| *Lactobacillus composti*_DSM_18527_JCM_14202 | *Paraburkholderia caryophylli*_Ballard_720 |
| *Peptoniphilus senegalensis*_JC140_type_strain_JC140 | *Marinospirillum minutulum*_DSM_6287 |
| *Pseudotherrmotoga thermarum*_DSM_5069 | *Halomonas cupida*_DSM_4740 |
| *Spiroplasma litorale*_TN_1 | *Mitsuokella multacida*_DSM_20544 |
| *Granulicella pectinivorans*_DSM_21001 | *Clostridium chauvoei*_JF4335 |
| *Poseidonocella sedimentorum*_KMM_9023_NRIC_0796_JCM_17311_KCTC_23692 | *Streptomyces lydicus*_103 |
| *Saccharomonospora xinjiangensis*_XJ_54 | *Xenorhabdus bovienii*_SS_2004 |
| *Rickettsia typhi*_str_Wilmington | *Rouxiella chamberiensis*_130333 |
| *Sulfitobacter brevis*_DSM_11443 | *Grimontia hollisae*_ATCC_33564 |
| *Treponema putidum*_OMZ_758 | *Veillonella parvula*_DSM_2008 |
| *Rickettsia japonica*_YH | *Robbsia andropogonis*_ICMP2807 |
| *Sporobacter termitidis*_DSM_10068 | *Pantoea agglomerans*_C410P1 |
| *Idiomarina salinarum*_ISL_52 | *Reichenbachiaella agariperforans*_DSM_26134 |
| *Clostridium hungatei*_DSM_14427 | *Selenomonas bovis*_DSM_23594 |
| *Clostridium papyrosolvens*_DSM_2782 | *Pacificimonas flava*_JLT2012 |
| *Nesterenkonia massiliensis*_NP1 | *Acinetobacter rudis*_DSM_24031 |
| *Thiothrix eikelboomii*_ATCC_49788 | *Legionella quinlivanii*_CDC_1442_AUS_E |
| *Leuconostoc mesenteroides*_subsp_mesenteroides_AT CC_8293 | *Burkholderia ubonensis*_MSMB22 |
| *Oceaniovalibus guishaninsula*_JLT2003 | *Carboxydotherrmus pertinax*_Ug1 |
| *Acetobacterium wieringae*_DSM_1911 | *Chryseobacterium formosense*_LMG_24722 |
| *Paracoccus zeaxanthinifaciens*_ATCC_21588 | *Aeromonas molluscorum*_848 |
| *Nocardia_miyunensis_NBRC_108239* | *Methylobacterium_mobilis_JLW8* |
| *Halolactibacillus_alkaliphilus_CGMCC_1_6843* | *Mycobacterium_wolinskyi_ATCC_700010* |
| *Olsenella_umbonata_KHGC19* | *Sediminibacterium_ginsengisoli_DSM_22335* |
| *Lactobacillus_paralimentarius_DSM_13961_JCM_10707* | *Halalkalibacillus_halophilus_DSM_18494* |
| *Sulfitobacter_noctilucae_NB_68* | *Edwardsiella_anguillarum_ET080813* |
| *Moorea_producens_PAL_8_15_08_1* | *Streptomyces_melanosporofaciens_DSM_40318* |
| *Helicobacter_macacae_MIT_99_5501* | *Stigmatella_aurantiaca_DW4_3_1* |
| *Pseudoalteromonas_spongiae_UST010723_006* | *Providencia_heimbachae_P12672* |
| *Acholeplasma_equifetale_ATCC_29724* | *Yersinia_pestis_CO92* |
| *Saccharibacter_floricola_DSM_15669* | *Nocardiopsis_trehalosi_NBRC_14201* |
| *Syntrophomonas_zehnderi_OL_4* | *Desulfuromusa_kysingii_DSM_7343* |
| *Oenococcus_kitaharae_DSM_17330* | *Halomonas_utahensis_P_halo* |
| *Thermus_brockianus_GE_1* | *Mycobacterium_colombiense_CECT_3035* |
| *Saccharomonospora_xinjiangensis_XJ_54* | *Marinobacter_nitratireducens_AK21* |
| *Tepidimonas_taiwanensis_VT154_175* | *Vibrio_vulnificus_CECT_4999* |
| *Faecalibacterium_prausnitzii_A2_165* | *Gracilibacillus_timonensis_Marseille_P2481* |
| *Alkalibacillus_haloalkaliphilus_C5* | *Clostridium_dakarensense_FF1* |
| *Lactobacillus_iners_DSM_13335* | *Aeromonas_hydrophila_subsp_hydrophila_ATCC_7966* |
| *Stenotrophomonas_ginsengisoli_DSM_24757* | *Xylophilus_ampelinus_CCH5_B3* |
| *Herbaspirillum_autotrophicum_IAM_14942* | *Algibacter_pectinivorans_DSM_25730* |
| *Treponema_caldarium_DSM_7334* | *Paraburkholderia_caryophylli_Ballard_720* |
| *Anaerostipes_caccae_DSM_14662* | *Pectobacterium_carotovorum_subsp_carotovorum_PC1* |
| *Lactobacillus_cacaonum_DSM_21116* | *Virgibacillus_necropolis_LMG_19488* |
| *Arcobacter_lanthieri_AF1440* | *Bacillus_testis_SIT10* |
| *Ruminococcus_flavifaciens_XPD3002* | *Dickeya_solani_IPO_2222* |
| *Desulfohalobium_retbaense_DSM_5692* | *Streptomyces_griseofuscus_NRRL_B_5429* |
| *Xanthomonas_maliensis_M97* | *Photobacterium_jeanii_R_40508* |
| *Dyadobacter_fermentans_DSM_18053* | *Bacillus_badius_SGD_V_25* |
| *Streptomyces_uncialis_DCA2648* | *Desulfonatronovibrio_magnus_AHT22* |
| *Lactobacillus_delbrueckii_subsp_bulgaricus_ATCC_11842_JCM_1002* | *Acinetobacter_soli_GFJ2* |
| *Streptobacillus_hongkongensis_DSM_26322* | *Streptomyces_guanduensis_CGMCC_4_2022* |
| *Acholeplasma_laidlawii_PG_8A* | *Pseudomonas_putida_PA14H7* |
| *Brachybacterium_massiliense_mt5* | *Prolixibacter_bellariivorans_ATCC_BAA_1284* |
| *Bacteroides_nordii_CL02T12C05* | *Aneurinibacillus_tyrosinisolvens_LL_002* |
| *Collinsella_stercoris_DSM_13279* | *Dethiosulfatarculus_sandiegensis_SPR* |
| *Leptospira_weilii_serovar_Topaz_str_LT2116* | *Lysobacter_enzymogenes_C3* |
| *Glutamicibacter_arilaitensis_KLBMP_5180* | *Xenorhabdus_japonica_DSM_16522* |
| *Nonomuraea_coxensis_DSM_45129* | *Tersicoccus_phoenicis_DSM_30849* |
| *Anaerovibrio_lipolyticus_DSM_3074* | *Deinococcus_wulumuqiensis_R13* |
| *Abiotrophia_defectiva_ATCC_49176* | *Bacteroides_pectinophilus_ATCC_43243* |
| *Desulfospira_joergensenii_DSM_10085* | *Vibrio_tubiaschii_ATCC_19109* |
| *Helicobacter_typhlonius* | *Moellerella_wisconsensis_ATCC_35017* |
| *Acidovorax_konjaci_DSM_7481* | *Halarsenatibacter_silvermanii_SLAS_1* |
| *Streptomyces_niveus_SCSIO_3406* | *Pseudomonas_fuscovaginae_LMG_2158* |
| *Lactobacillus_senioris_DSM_24302_JCM_17472* | *Nakamurella_lactea_DSM_19367* |
| *Aerococcus_urinaehominis_CCUG42038B* | *Xenorhabdus_bovienii_SS_2004* |
| *Pseudarcicella_hirudinis_E92_LMG_26720_CCM_7988* | *Nocardiopsis_potens_DSM_45234* |
| *Desulfurobacterium_indicum_K6013* | *Halomonas_elongata_HEK1* |
| *Massiliomicrobiota_timonensis_SN16* | *Citrobacter_freundii_CFNH1* |
| *Nocardiopsis_salina_YIM_90010* | *Gulbenkiania_indica_DSM_17901* |
| *Streptococcus_sanguinis_SK36* | *Endozoicomonas_elysicola_DSM_22380* |
| *Helicobacter_macacae_MIT_99_5501* | *Nocardiopsis_xinjiangensis_YIM_90004* |

**Table S8.**
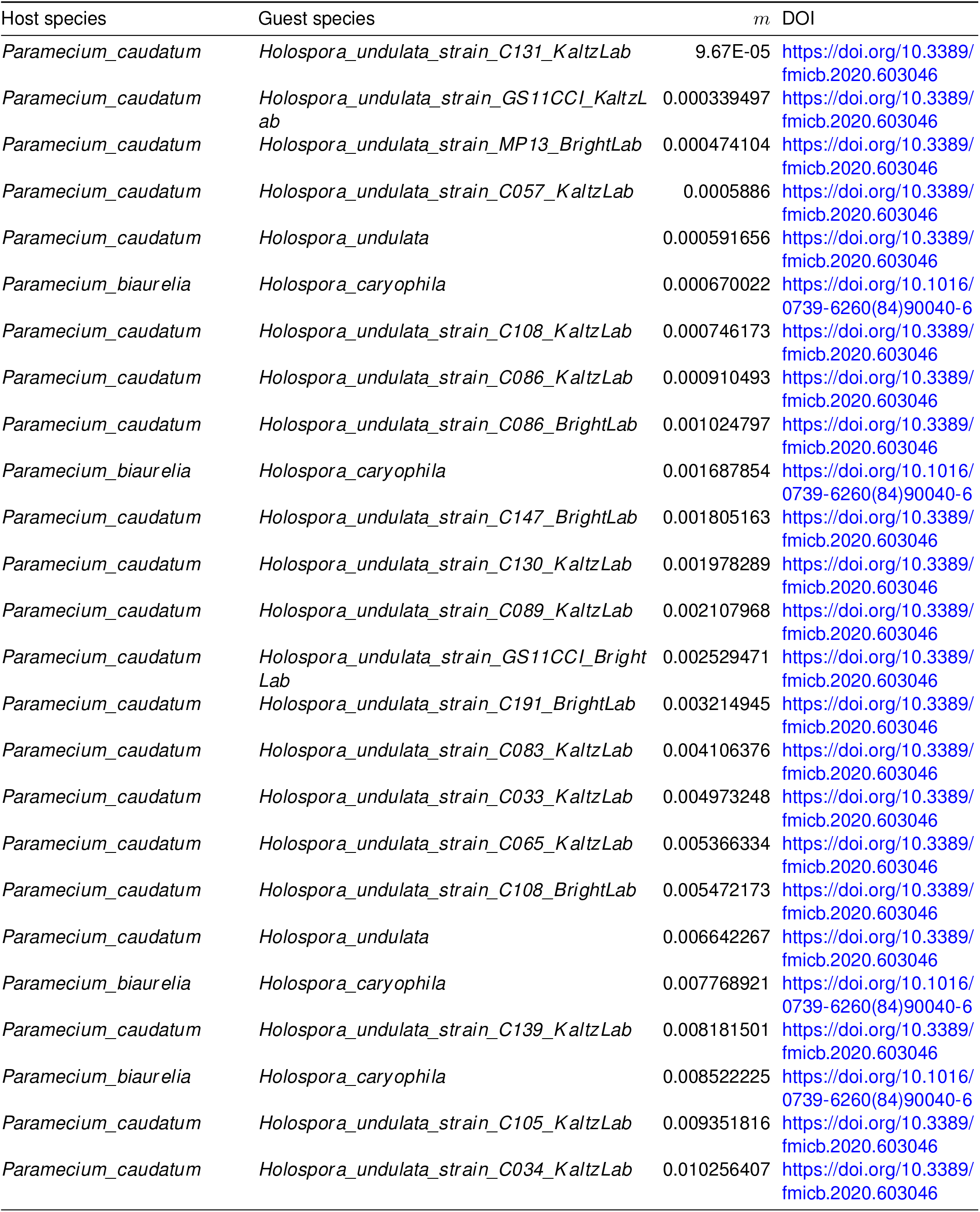
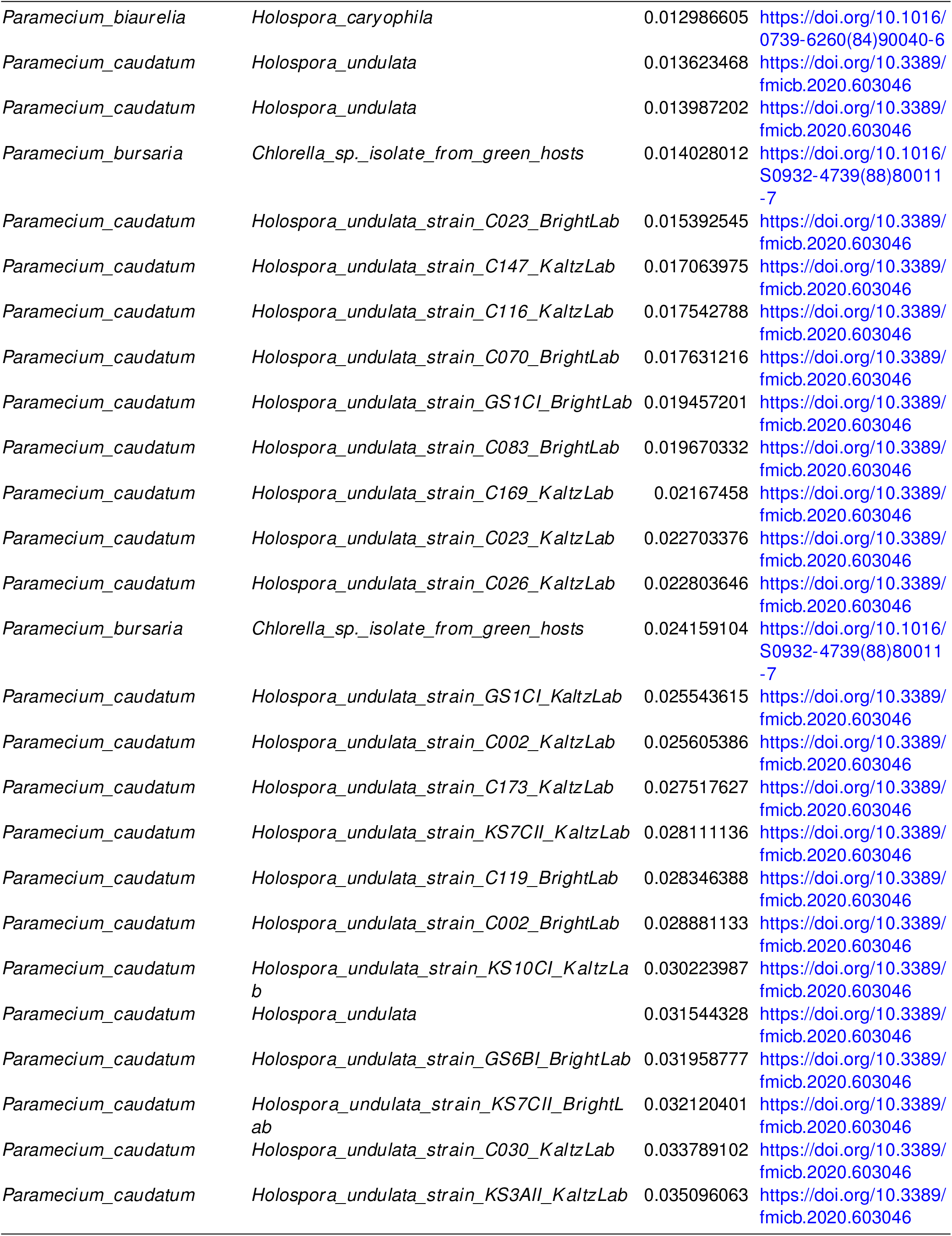
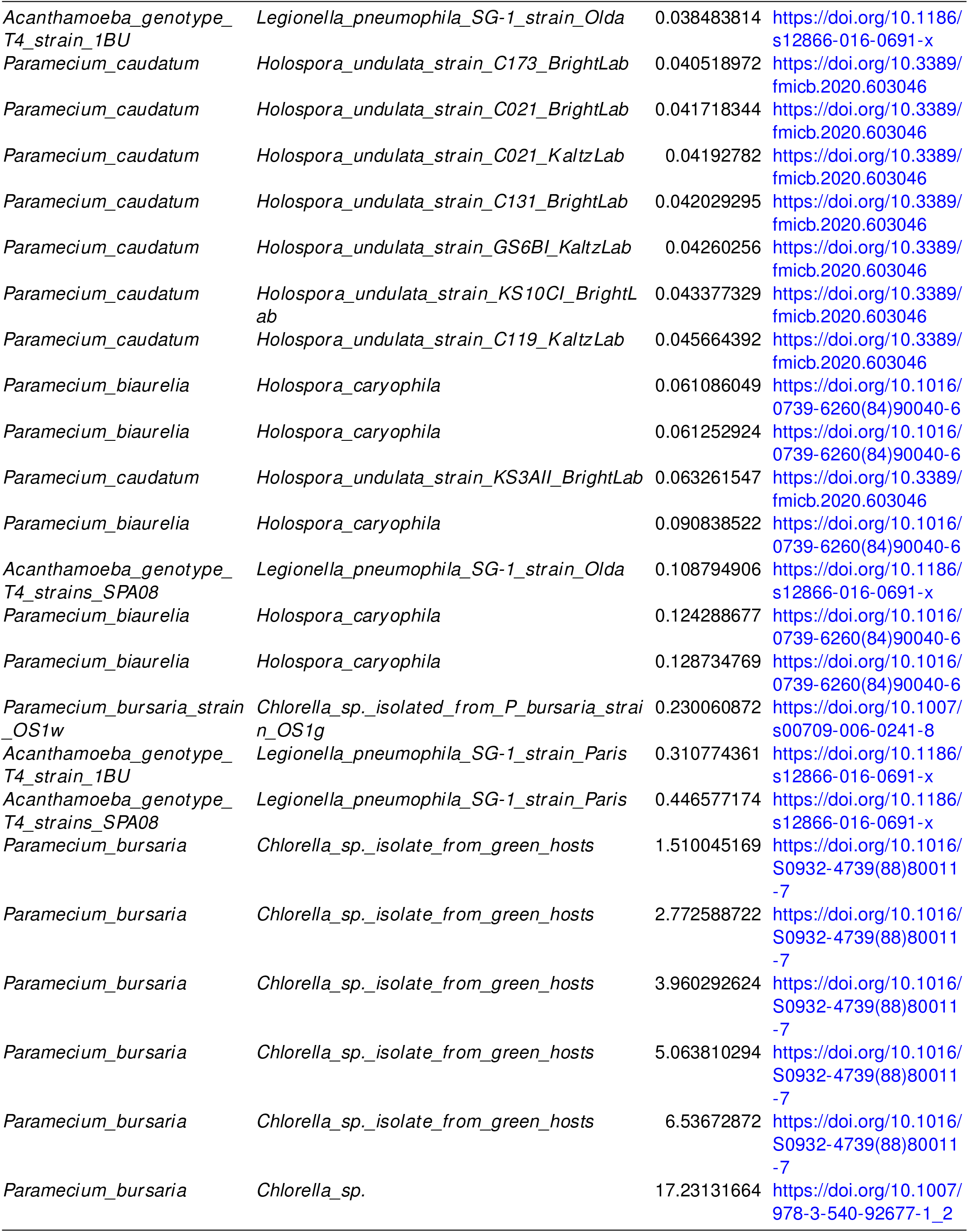
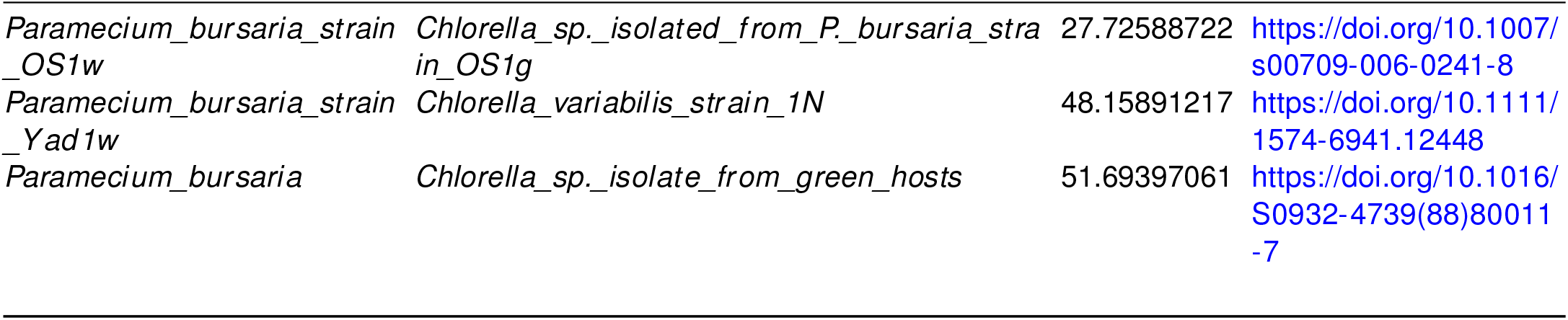
This table summarizes data from eight studies (45–51), reporting the host and guest species and the frequency *f* of initially guest-free host populations that harbored guests after a time *T*. From these reported values of *f* and *T*, we estimated the per-capita rate of new endosymbiosis formation as *m* = − ln(1 −*f*)/*T*. The resulting estimates are plotted in Fig. 4D and Fig. S3B.

| Host species | Guest species | $m$ | DOI |
| --- | --- | --- | --- |
| *Paramecium_caudatum* | *Holospira_undulata_strain_C131_KaltzLab* | 9.67E-05 | https://doi.org/10.3389/fmicb.2020.603046 |
| *Paramecium_caudatum* | *Holospira_undulata_strain_GS11CCI_KaltzLab* | 0.000339497 | https://doi.org/10.3389/fmicb.2020.603046 |
| *Paramecium_caudatum* | *Holospira_undulata_strain_MP13_BrightLab* | 0.000474104 | https://doi.org/10.3389/fmicb.2020.603046 |
| *Paramecium_caudatum* | *Holospira_undulata_strain_C057_KaltzLab* | 0.0005886 | https://doi.org/10.3389/fmicb.2020.603046 |
| *Paramecium_caudatum* | *Holospira_undulata* | 0.000591656 | https://doi.org/10.3389/fmicb.2020.603046 |
| *Paramecium_biaurelia* | *Holospira_caryophila* | 0.000670022 | [https://doi.org/10.1016/0739-6260(84)90040-6](https://doi.org/10.1016/0739-6260%2884%2990040-6) |
| *Paramecium_caudatum* | *Holospira_undulata_strain_C108_KaltzLab* | 0.000746173 | https://doi.org/10.3389/fmicb.2020.603046 |
| *Paramecium_caudatum* | *Holospira_undulata_strain_C086_KaltzLab* | 0.000910493 | https://doi.org/10.3389/fmicb.2020.603046 |
| *Paramecium_caudatum* | *Holospira_undulata_strain_C086_BrightLab* | 0.001024797 | https://doi.org/10.3389/fmicb.2020.603046 |
| *Paramecium_biaurelia* | *Holospira_caryophila* | 0.001687854 | [https://doi.org/10.1016/0739-6260(84)90040-6](https://doi.org/10.1016/0739-6260%2884%2990040-6) |
| *Paramecium_caudatum* | *Holospira_undulata_strain_C147_BrightLab* | 0.001805163 | https://doi.org/10.3389/fmicb.2020.603046 |
| *Paramecium_caudatum* | *Holospira_undulata_strain_C130_KaltzLab* | 0.001978289 | https://doi.org/10.3389/fmicb.2020.603046 |
| *Paramecium_caudatum* | *Holospira_undulata_strain_C089_KaltzLab* | 0.002107968 | https://doi.org/10.3389/fmicb.2020.603046 |
| *Paramecium_caudatum* | *Holospira_undulata_strain_GS11CCI_BrightLab* | 0.002529471 | https://doi.org/10.3389/fmicb.2020.603046 |
| *Paramecium_caudatum* | *Holospira_undulata_strain_C191_BrightLab* | 0.003214945 | https://doi.org/10.3389/fmicb.2020.603046 |
| *Paramecium_caudatum* | *Holospira_undulata_strain_C083_KaltzLab* | 0.004106376 | https://doi.org/10.3389/fmicb.2020.603046 |
| *Paramecium_caudatum* | *Holospira_undulata_strain_C033_KaltzLab* | 0.004973248 | https://doi.org/10.3389/fmicb.2020.603046 |
| *Paramecium_caudatum* | *Holospira_undulata_strain_C065_KaltzLab* | 0.005366334 | https://doi.org/10.3389/fmicb.2020.603046 |
| *Paramecium_caudatum* | *Holospira_undulata_strain_C108_BrightLab* | 0.005472173 | https://doi.org/10.3389/fmicb.2020.603046 |
| *Paramecium_caudatum* | *Holospira_undulata* | 0.006642267 | https://doi.org/10.3389/fmicb.2020.603046 |
| *Paramecium_biaurelia* | *Holospira_caryophila* | 0.007768921 | [https://doi.org/10.1016/0739-6260(84)90040-6](https://doi.org/10.1016/0739-6260%2884%2990040-6) |
| *Paramecium_caudatum* | *Holospira_undulata_strain_C139_KaltzLab* | 0.008181501 | https://doi.org/10.3389/fmicb.2020.603046 |
| *Paramecium_biaurelia* | *Holospira_caryophila* | 0.008522225 | [https://doi.org/10.1016/0739-6260(84)90040-6](https://doi.org/10.1016/0739-6260%2884%2990040-6) |
| *Paramecium_caudatum* | *Holospira_undulata_strain_C105_KaltzLab* | 0.009351816 | https://doi.org/10.3389/fmicb.2020.603046 |
| *Paramecium_caudatum* | *Holospira_undulata_strain_C034_KaltzLab* | 0.010256407 | https://doi.org/10.3389/fmicb.2020.603046 |

| Host species | Guest species | *m* | DOI |
| --- | --- | --- | --- |
| *Paramecium_biaurelia* | *Holospora_caryophila* | 0.012986605 | [https://doi.org/10.1016/0739-6260(84)90040-6](https://doi.org/10.1016/0739-6260%2884%2990040-6) |
| *Paramecium_caudatum* | *Holospora_undulata* | 0.013623468 | https://doi.org/10.3389/fmicb.2020.603046 |
| *Paramecium_caudatum* | *Holospora_undulata* | 0.013987202 | https://doi.org/10.3389/fmicb.2020.603046 |
| *Paramecium_bursaria* | *Chlorella_sp._isolate_from_green_hosts* | 0.014028012 | [https://doi.org/10.1016/S0932-4739(88)80011-7](https://doi.org/10.1016/S0932-4739%2888%2980011-7) |
| *Paramecium_caudatum* | *Holospora_undulata_strain_C023_BrightLab* | 0.015392545 | https://doi.org/10.3389/fmicb.2020.603046 |
| *Paramecium_caudatum* | *Holospora_undulata_strain_C147_KaltzLab* | 0.017063975 | https://doi.org/10.3389/fmicb.2020.603046 |
| *Paramecium_caudatum* | *Holospora_undulata_strain_C116_KaltzLab* | 0.017542788 | https://doi.org/10.3389/fmicb.2020.603046 |
| *Paramecium_caudatum* | *Holospora_undulata_strain_C070_BrightLab* | 0.017631216 | https://doi.org/10.3389/fmicb.2020.603046 |
| *Paramecium_caudatum* | *Holospora_undulata_strain_GS1CI_BrightLab* | 0.019457201 | https://doi.org/10.3389/fmicb.2020.603046 |
| *Paramecium_caudatum* | *Holospora_undulata_strain_C083_BrightLab* | 0.019670332 | https://doi.org/10.3389/fmicb.2020.603046 |
| *Paramecium_caudatum* | *Holospora_undulata_strain_C169_KaltzLab* | 0.02167458 | https://doi.org/10.3389/fmicb.2020.603046 |
| *Paramecium_caudatum* | *Holospora_undulata_strain_C023_KaltzLab* | 0.022703376 | https://doi.org/10.3389/fmicb.2020.603046 |
| *Paramecium_caudatum* | *Holospora_undulata_strain_C026_KaltzLab* | 0.022803646 | https://doi.org/10.3389/fmicb.2020.603046 |
| *Paramecium_bursaria* | *Chlorella_sp._isolate_from_green_hosts* | 0.024159104 | [https://doi.org/10.1016/S0932-4739(88)80011-7](https://doi.org/10.1016/S0932-4739%2888%2980011-7) |
| *Paramecium_caudatum* | *Holospora_undulata_strain_GS1CI_KaltzLab* | 0.025543615 | https://doi.org/10.3389/fmicb.2020.603046 |
| *Paramecium_caudatum* | *Holospora_undulata_strain_C002_KaltzLab* | 0.025605386 | https://doi.org/10.3389/fmicb.2020.603046 |
| *Paramecium_caudatum* | *Holospora_undulata_strain_C173_KaltzLab* | 0.027517627 | https://doi.org/10.3389/fmicb.2020.603046 |
| *Paramecium_caudatum* | *Holospora_undulata_strain_KS7CII_KaltzLab* | 0.028111136 | https://doi.org/10.3389/fmicb.2020.603046 |
| *Paramecium_caudatum* | *Holospora_undulata_strain_C119_BrightLab* | 0.028346388 | https://doi.org/10.3389/fmicb.2020.603046 |
| *Paramecium_caudatum* | *Holospora_undulata_strain_C002_BrightLab* | 0.028881133 | https://doi.org/10.3389/fmicb.2020.603046 |
| *Paramecium_caudatum* | *Holospora_undulata_strain_KS10CI_KaltzLab* | 0.030223987 | https://doi.org/10.3389/fmicb.2020.603046 |
| *Paramecium_caudatum* | *Holospora_undulata* | 0.031544328 | https://doi.org/10.3389/fmicb.2020.603046 |
| *Paramecium_caudatum* | *Holospora_undulata_strain_GS6BI_BrightLab* | 0.031958777 | https://doi.org/10.3389/fmicb.2020.603046 |
| *Paramecium_caudatum* | *Holospora_undulata_strain_KS7CII_BrightLab* | 0.032120401 | https://doi.org/10.3389/fmicb.2020.603046 |
| *Paramecium_caudatum* | *Holospora_undulata_strain_C030_KaltzLab* | 0.033789102 | https://doi.org/10.3389/fmicb.2020.603046 |
| *Paramecium_caudatum* | *Holospora_undulata_strain_KS3All_KaltzLab* | 0.035096063 | https://doi.org/10.3389/fmicb.2020.603046 |

| Host species | Guest species | *m* | DOI |
| --- | --- | --- | --- |
| *Acanthamoeba*_genotype_T4_strain_1BU | *Legionella_pneumophila*_SG-1_strain_Old | 0.038483814 | https://doi.org/10.1186/s12866-016-0691-x |
| *Paramecium*_caudatum | *Holospora_undulata*_strain_C173_BrightLab | 0.040518972 | https://doi.org/10.3389/fmicb.2020.603046 |
| *Paramecium*_caudatum | *Holospora_undulata*_strain_C021_BrightLab | 0.041718344 | https://doi.org/10.3389/fmicb.2020.603046 |
| *Paramecium*_caudatum | *Holospora_undulata*_strain_C021_KaltzLab | 0.04192782 | https://doi.org/10.3389/fmicb.2020.603046 |
| *Paramecium*_caudatum | *Holospora_undulata*_strain_C131_BrightLab | 0.042029295 | https://doi.org/10.3389/fmicb.2020.603046 |
| *Paramecium*_caudatum | *Holospora_undulata*_strain_GS6BI_KaltzLab | 0.04260256 | https://doi.org/10.3389/fmicb.2020.603046 |
| *Paramecium*_caudatum | *Holospora_undulata*_strain_KS10CI_BrightLab | 0.043377329 | https://doi.org/10.3389/fmicb.2020.603046 |
| *Paramecium*_caudatum | *Holospora_undulata*_strain_C119_KaltzLab | 0.045664392 | https://doi.org/10.3389/fmicb.2020.603046 |
| *Paramecium_biaurelia* | *Holospora_caryophila* | 0.061086049 | [https://doi.org/10.1016/0739-6260(84)90040-6](https://doi.org/10.1016/0739-6260%2884%2990040-6) |
| *Paramecium_biaurelia* | *Holospora_caryophila* | 0.061252924 | [https://doi.org/10.1016/0739-6260(84)90040-6](https://doi.org/10.1016/0739-6260%2884%2990040-6) |
| *Paramecium*_caudatum | *Holospora_undulata*_strain_KS3All_BrightLab | 0.063261547 | https://doi.org/10.3389/fmicb.2020.603046 |
| *Paramecium_biaurelia* | *Holospora_caryophila* | 0.090838522 | [https://doi.org/10.1016/0739-6260(84)90040-6](https://doi.org/10.1016/0739-6260%2884%2990040-6) |
| *Acanthamoeba*_genotype_T4_strains_SPA08 | *Legionella_pneumophila*_SG-1_strain_Old | 0.108794906 | https://doi.org/10.1186/s12866-016-0691-x |
| *Paramecium_biaurelia* | *Holospora_caryophila* | 0.124288677 | [https://doi.org/10.1016/0739-6260(84)90040-6](https://doi.org/10.1016/0739-6260%2884%2990040-6) |
| *Paramecium_biaurelia* | *Holospora_caryophila* | 0.128734769 | [https://doi.org/10.1016/0739-6260(84)90040-6](https://doi.org/10.1016/0739-6260%2884%2990040-6) |
| *Paramecium_bursaria*_strain_OS1w | *Chlorella_sp._isolated_from_P_bursaria_strain_OS1g* | 0.230060872 | https://doi.org/10.1007/s00709-006-0241-8 |
| *Acanthamoeba*_genotype_T4_strain_1BU | *Legionella_pneumophila*_SG-1_strain_Paris | 0.310774361 | https://doi.org/10.1186/s12866-016-0691-x |
| *Acanthamoeba*_genotype_T4_strains_SPA08 | *Legionella_pneumophila*_SG-1_strain_Paris | 0.446577174 | https://doi.org/10.1186/s12866-016-0691-x |
| *Paramecium_bursaria* | *Chlorella_sp._isolate_from_green_hosts* | 1.510045169 | [https://doi.org/10.1016/S0932-4739(88)80011-7](https://doi.org/10.1016/S0932-4739%2888%2980011-7) |
| *Paramecium_bursaria* | *Chlorella_sp._isolate_from_green_hosts* | 2.772588722 | [https://doi.org/10.1016/S0932-4739(88)80011-7](https://doi.org/10.1016/S0932-4739%2888%2980011-7) |
| *Paramecium_bursaria* | *Chlorella_sp._isolate_from_green_hosts* | 3.960292624 | [https://doi.org/10.1016/S0932-4739(88)80011-7](https://doi.org/10.1016/S0932-4739%2888%2980011-7) |
| *Paramecium_bursaria* | *Chlorella_sp._isolate_from_green_hosts* | 5.063810294 | [https://doi.org/10.1016/S0932-4739(88)80011-7](https://doi.org/10.1016/S0932-4739%2888%2980011-7) |
| *Paramecium_bursaria* | *Chlorella_sp._isolate_from_green_hosts* | 6.53672872 | [https://doi.org/10.1016/S0932-4739(88)80011-7](https://doi.org/10.1016/S0932-4739%2888%2980011-7) |
| *Paramecium_bursaria* | *Chlorella_sp.* | 17.23131664 | https://doi.org/10.1007/978-3-540-92677-1_2 |

| Host species | Guest species | *m* | DOI |
| --- | --- | --- | --- |
| *Paramecium_bursaria_strain_OS1w* | *Chlorella_sp._isolated_from_P._bursaria_strain_OS1g* | 27.72588722 | https://doi.org/10.1007/s00709-006-0241-8 |
| *Paramecium_bursaria_strain_Yad1w* | *Chlorella_variabilis_strain_1N* | 48.15891217 | https://doi.org/10.1111/1574-6941.12448 |
| *Paramecium_bursaria* | *Chlorella_sp._isolate_from_green_hosts* | 51.69397061 | [https://doi.org/10.1016/S0932-4739(88)80011-7](https://doi.org/10.1016/S0932-4739%2888%2980011-7) |

### A. Derivation of dissociation rate

To get an estimate of the dissociation rate, we develop a simple mathematical model. Suppose there is a host-guest pair *H*_1_ that grows according to some exponential rate *k* but loses its guest at rate *d*. We can write the dynamics of this population as the differential equation:

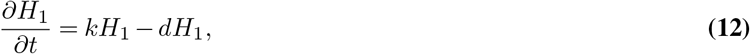

which means the population *H*_1_ grows exponentially according to

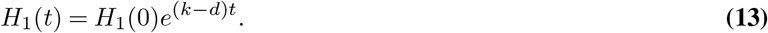

This model assumes continuous population growth, but we can also express growth in terms of discrete reproductive events in which populations double at some characteristic time *τ*. In this case, a population without dissociation would grow as *H*_1_(*t*) = *H*_1_(0)2^*t/τ*^. We can include dissociation by assuming that with probability *p* one daughter cell will not get the guest (note that at least one daughter is guaranteed to get the guest). With dissociation, the expression for population growth becomes *H*_1_(*t*) = *H*_1_(0)(1 + 1 −*p*)^*t/τ*^ or equivalently

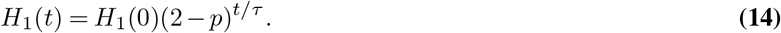

The *p* term is the probability that upon reproduction the guest is not transmitted to a host offspring, and *τ* is the characteristic reproductive time (*τ* = ln 2/*k*). This alternative formulation has the benefit that it recasts dissociation in terms of a probability rather than a rate, which is easier to estimate. By setting the two formulations for population growth equal, we can write the dissociation rate *d* in terms of *p* and *k*:

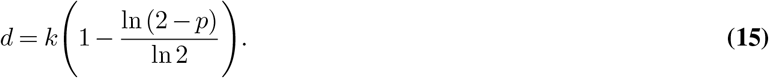

Since *k* is the maximum growth rate of hosts, we can estimate it with the metabolic models, i.e. *k* is *µ*_*u*_, which is proportional to *W*_*H*_. Thus, all we need to determine *d* is to estimate *p*.

We can estimate *p* by using the ratio of the doubling times. In order for the host-guest pair to successfully be replicated, the guest needs to reproduce by the time the host reproduces. So if the host’s doubling time is *τ*_*h*_, the guest needs to reproduce within this window. If the guest takes longer than this time, then it will miss one window, but it might make the next. This would count as one failure and one success in terms of host-guest reproduction for a value *p* = 0.5. A simple way to compute *p* is to ask within a given time interval how many times the guest reproduced and how many times the host reproduced. For example, if *τ*_*h*_ = 1 and *τ*_*e*_ = 1.4, then in some interval *t* the host reproduces *t/τ*_*h*_ times and the guest *t/τ*_*e*_ times. If these two values are the same, then there would be no dissociation events, which would mean *p* = 0. More generally, we can write the value of *p* as:

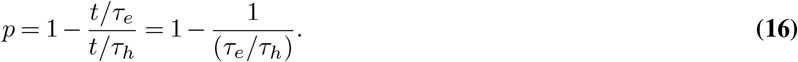

We can use the relationship between doubling times *τ* and growth rates of populations *k* to put the ratio of doubling times in terms of the ratio of growth rates.

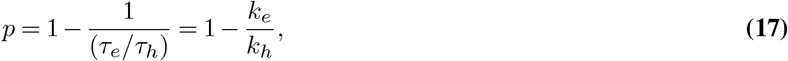

where *k*_*e*_ is the growth rate of the guest in the endosymbiosis and *k*_*h*_ is the growth rate of the host in the endosymbiosis. Lastly, by substituting Eq. (17) into Eq. (15), we have that

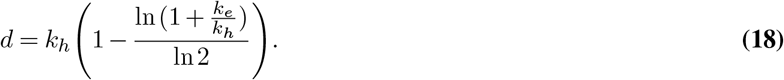

We note that in the main text the rate *k*_*h*_ is equivalent to *µ*_*u*_ and *k*_*e*_ is equivalent to *µ*_*g*_.

### B. Calculating the per-host guest acquisition rate from literature data

In most empirical studies, authors report the fraction *f* of hosts that have acquired a guest after a fixed observation time *T*. To compare this data with our mathematical modeling, we need to convert those fractions into terms of per-capita conversion of hosts without guests into hosts carrying guests, i.e. *m*. The short derivation below converts the reported fraction *f* into the rate *m*.

Before presenting a simple mathematical model, we make the following three assumptions. First, we assume that the total host population is not growing, so that the sum of hosts with guests and hosts without guests is constant. Second, we assume that the concentration of free-living guests is sufficiently high that the formation rate of endosymbioses is effectively constant. This assumption is equivalent to assuming that there are many more guests than hosts. Finally, we assume that each host can only acquire one guest. A consequence of these assumptions is that our model will underestimate the actual rate *m* in the empirical systems. However, our argument in the manuscript is that our calculated rates of *m*\* are not so high as to be unreasonable. Thus, if the *m* in real systems is higher than our estimate then that only serves to support our claim.

Based on our assumptions, we can write the proportion *P* (*t*) of hosts that are without guests at time *t* as

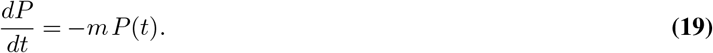

Here, *m* is the rate that hosts acquire guests. We note that *P* (*t*) is equivalent to the ratio of the number/concentration of hosts without guests divided by the total number/concentration of hosts in a population. The empirical studies typically begin with a uniform host population that does not have guests, and so *P* (0) = 1. We solve for *P* (*t*) to get

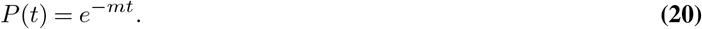

After an observation period *T*, empirical studies measure the fraction of the population with guests. This fraction *f* is simply:

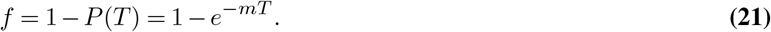

Solving for *m* gives the following equation:

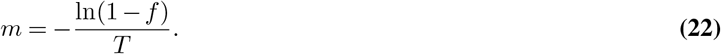

### C. Comparison of *m*\* and *d*

We want to compare the values of *m*\* and *d* in the differential equations model using data computed from the metabolic models. The growth rates in the metabolic models and the growth rates in the differential equations models are not in the same units. However, there are ratios of growth rates in the two models that are equivalent, e.g. *µ*_*s*_/*µ*_*u*_ = *W*_*S*_/*W*_*H*_. We note that we can rewrite the expression for *d* in Eq. (2) as:

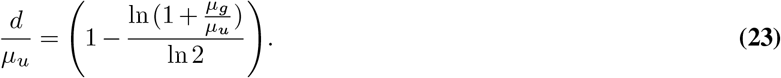

We can also rewrite the expression for *m*\* found in Eq. (5) as:

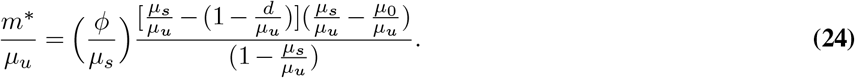

The benefit of this reformulation is that the expressions for 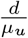 and 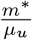 are in terms of ratios 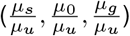 that are equivalent to ratios that can be computed using metabolic models. We also note that this rescaling of *d* and *m*\* by *µ*_*u*_ can be found by rescaling time *t* in the differential equations. By letting *τ* = *µ*_*u*_*t* we can rewrite Eq. (4) as:

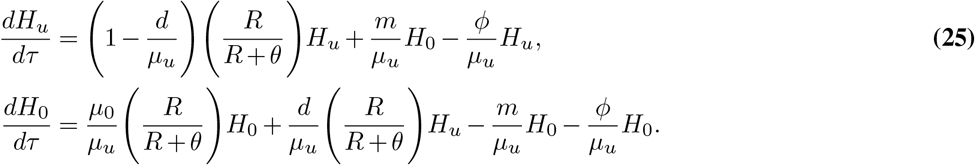

The only terms unaccounted for can be related by the following equivalence: 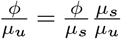.

### D. 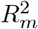calculation

To quantify the proportion of variance of *m*\*/*µ*_*u*_ explained by *d*/*µ*_*u*_, we fitted a linear mixed-effects model with crossed random intercepts for guests and hosts. This formulation accounts for the non-independence: although each observation comes from a unique host-guest pair, the same host or guest may appear in multiple pairs. This data structure induces correlation among observations that share an individual, which, if ignored, can inflate *R*^2^ by wrongly attributing variance due to shared host/guest baselines to the fixed effect.

Let *y*_*gh*_ = *m*\*/*µ*_*u*_ be the response for the pair formed by guest *g* with host *h*, and *x*_*gh*_ = *d/µ*_*u*_ be the explanatory variable. The model is

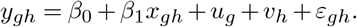

Here, *g* ∈ {1,…, *G*} indexes guests and *h* ∈ {1,…, *H*} indexes hosts. For AGORA, *G* = 787 and *H* = 696; for CarveMe, *G* = 5044 and *H* = 5182. The random terms *u*_*g*_ and *v*_*h*_ represent guest- and host-specific deviations from the intercept, with 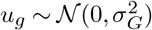, and 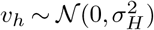. The residual 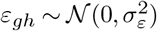 captures observation-level variability not explained by the fixed effect or the random intercepts.

We quantified *R*^2^ marginal 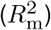 as the proportion of variance explained by the fixed effect:

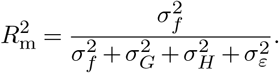

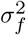 is the variance across observations of the fixed-effect predictor. We found 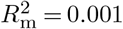 for AGORA and 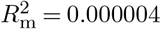 for CarveMe. However, we found nonlinearity and the presence of influential points, suggesting that our calculated 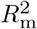 might be inaccurate. To address this issue, we re-expressed the model using the underlying theoretical relationship between *m*\*/*µ*_*u*_ and *d/µ*_*u*_. Specifically, this relationship originates from the equation:

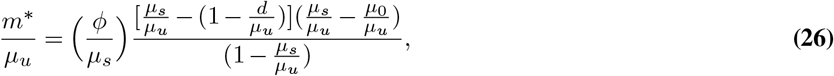

which we rewrote in the equivalent mathematical formulation:

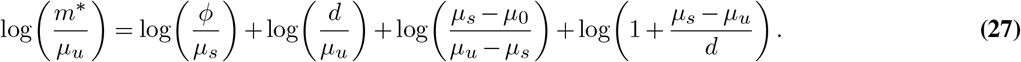

Using the same mixed-effects structure with *y*_*gh*_ = log_10_(*m*\*/*µ*_*u*_) and *x*_*gh*_ = log_10_(*d/µ*_*u*_), we found linearity and only weak evidence of influential points, with 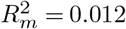 (AGORA) and 0.003 (CarveMe).

We ran the linear mixed-effects model using the `lmer` function, and calculated 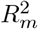 with the `r2_nakagawa` function in R version 4.4.3.

